# The recount3 Python package for programmatic access to uniformly processed RNA-seq data

**DOI:** 10.64898/2026.06.17.732943

**Authors:** Alexander Alsalihi, Robert M. Flight, Hunter N. B. Moseley

## Abstract

The recount3 online resource provides tens of thousands of uniformly processed RNA-seq samples across human and mouse from major sequencing repositories like the Sequence Read Archive. While access to these datasets has traditionally been centered in the R/Bioconductor ecosystem, the growing prominence of Python in bioinformatics and machine learning necessitates native, efficient tooling for Python users. Therefore, we present the recount3 Python package with robust application programming interface (API) and command-line interface (CLI) for discovering, downloading, and materializing recount3 resources. The software orchestrates uniform resource locator (URL) resolution, persistent on-disk caching, and the automatic parsing of data into analysis-ready data structures, including Pandas DataFrames and BiocPy RangedSummarizedExperiment objects. The recount3 Python package drastically lowers the barrier to entry for large-scale utilization of RNA-seq data in Python-based computational pipelines, bridging the gap between massive public transcriptomic data and modern machine learning ecosystems.

## Background

For roughly twenty years, next generation sequencing platforms using the RNA-seq technique [1] have generated a wealth of gene expression data, which has been deposited in sequencing repositories like the Sequence Read Archive (SRA) [2]. While this wealth of gene expression data has been a boon to biomedical research, large-scale reuse of this data has been limited to very few groups with high transcriptomics data analysis expertise and large computational resources. Uniformly processing a large number of raw RNA-seq datasets from the SRA is a daunting task for most biomedical researchers, even researchers with substantial transcriptomics data analysis expertise. To lower this reuse burden, the recount3 online resource was created as a derivative repository of uniformly-processed human and mouse RNA-seq datasets [3]. This resource utilizes raw RNA-seq datasets from major sequencing repositories including the SRA, the Genotype-Tissue Expression (GTEx) Portal [4], and The Cancer Genome Atlas (TCGA) [5]. The recount3 online resource provides tens of thousands of uniformly-processed RNA-seq datasets, enabling large-scale reuse of biomedically-relevant gene expression data with minimal reprocessing. The recount3 and snapcount R/Bioconductor packages provide R programmatic access to this online resource, especially for differential analyses [3]. However, access to recount3 in the Python programming language or more generally on the command line has been lacking until now. Here, we present the recount3 Python package which provides both a Python application programming interface (API) and a command line interface (CLI) for general access to the recount3 online resource.

## Results

The recount3 Python package supports data exploration and bioinformatics workflows through an API structured into three layers of abstraction: high-level experiment builders, mid-level resource bundles, and low-level resource managers. The following use cases demonstrate the package’s core functionalities.

### High-level dataset assembly

The create_rse() function provides the highest-level entry point for dataset assembly. It automates resource discovery, downloading, metadata alignment, and data parsing. By supplying a project identifier, organism, and annotation label, users query the repository to construct a RangedSummarizedExperiment (RSE) object. This BiocPy-compatible object integrates the feature-by-sample count matrix, sample metadata, and genomic coordinates into a single data structure. A single call performs every step illustrated in Figure 3:

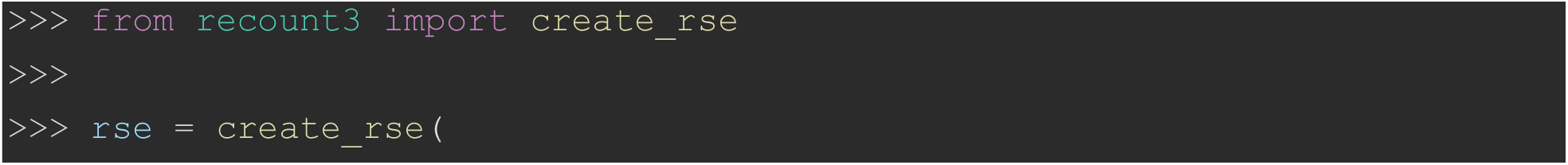
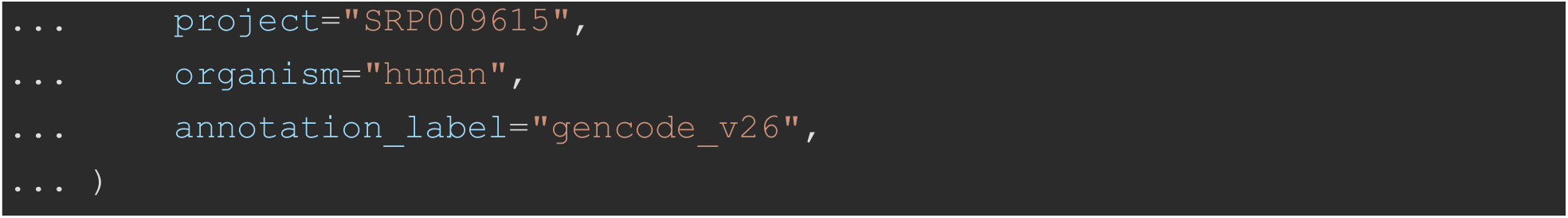

The function supports gene-, exon-, and junction-level summarizations via the genomic_unit argument. For junction-level assemblies, the function automatically retrieves and parses the “RR” (row ranges) sidecar file to attach genomic coordinates to each junction row:

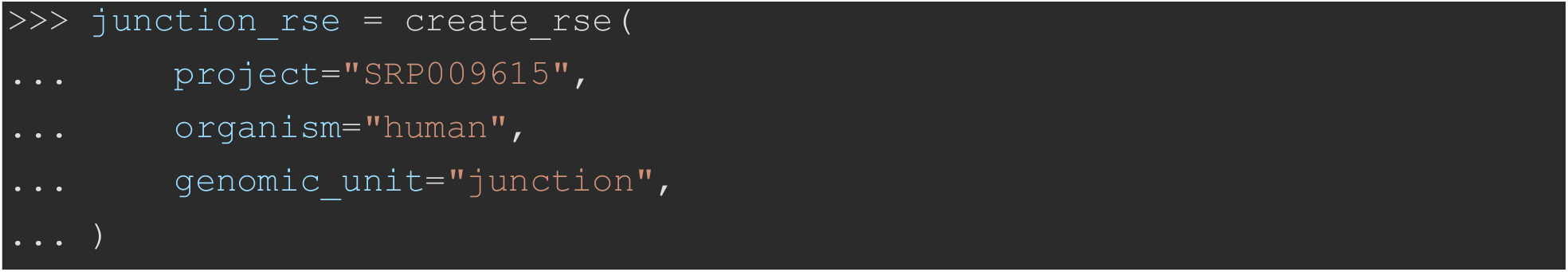

Annotation builds are addressed either by human-readable label (annotation_label=“gencode_v26”) or by the underlying extension code (annotation_extension=“G026”). The explicit extension takes precedence when both are supplied. The full list of supported labels for a given organism can be inspected programmatically:

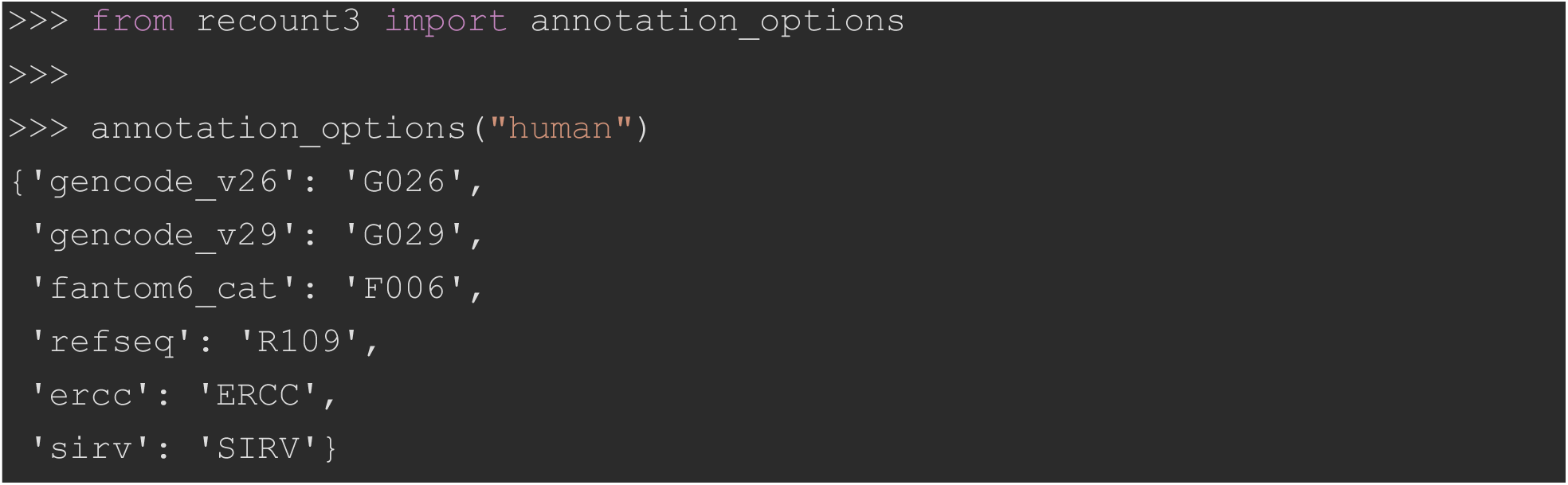

Mouse builds are exposed the same way:

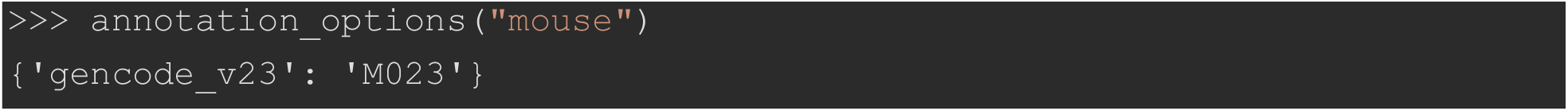

If genomic ranges are unavailable for a given parameter combination, users can fall back to a plain SummarizedExperiment object by setting allow_fallback_to_se=True:

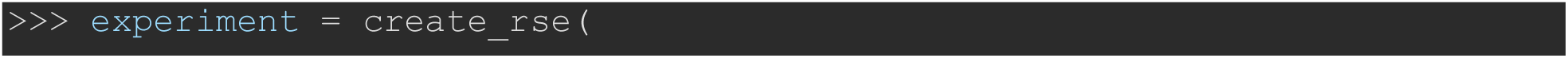
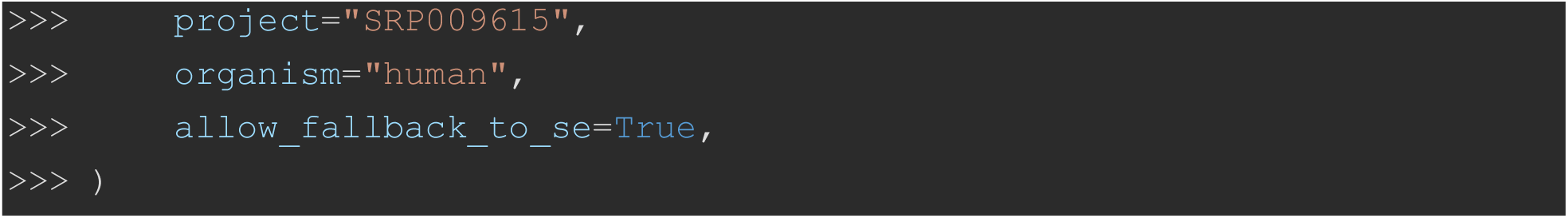

### Mid-level discovery and multi-project aggregation

For analyses spanning multiple studies or requiring data subsetting prior to materialization, the mid-level R3ResourceBundle class is used. The discover() class method accepts iterable objects, such as lists of project identifiers or organisms, and computes the Cartesian product to locate all relevant files, returning a unified resource bundle:

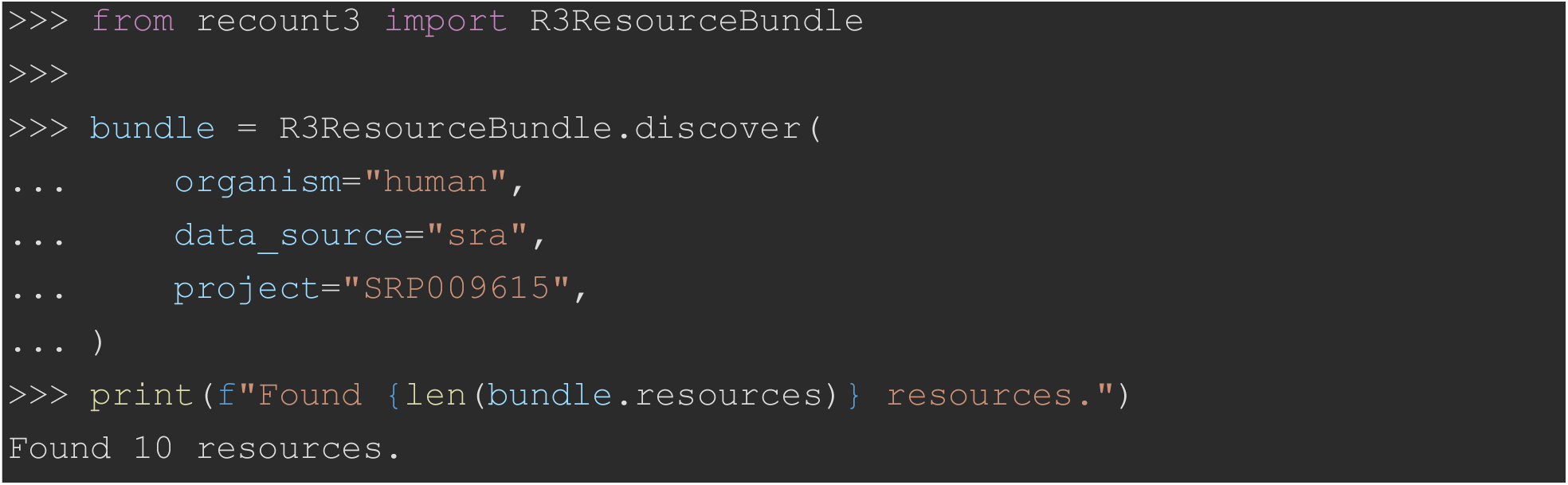

By providing the scalar project argument with a list, the discover class method aggregates every matching resource into a single bundle, which preserves per-resource identity while omitting bundle-level identity attributes to avoid misrepresentation:

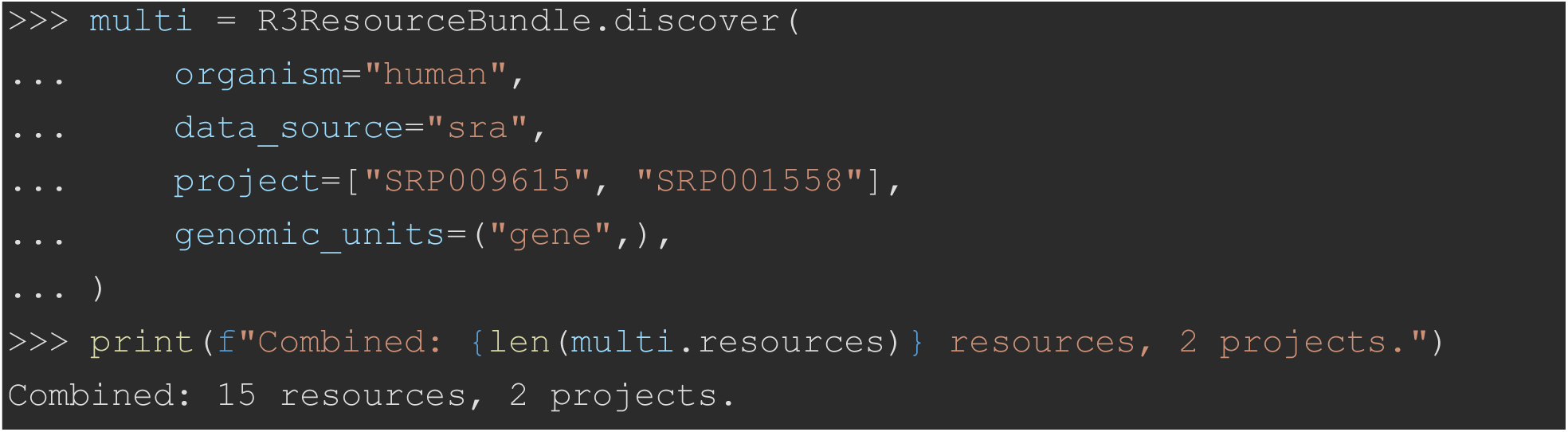

Users can subset bundles using the filter() method, which accepts exact strings, iterables of strings, or custom callable predicates against any field of the underlying resource description. Filtering results are reversed with the invert=True parameter option. The following idioms cover the most common selection patterns:

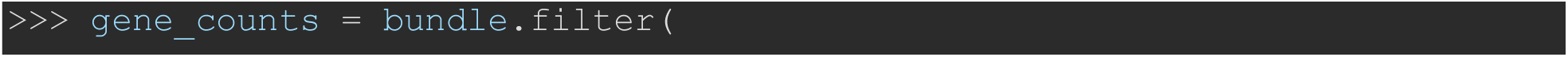
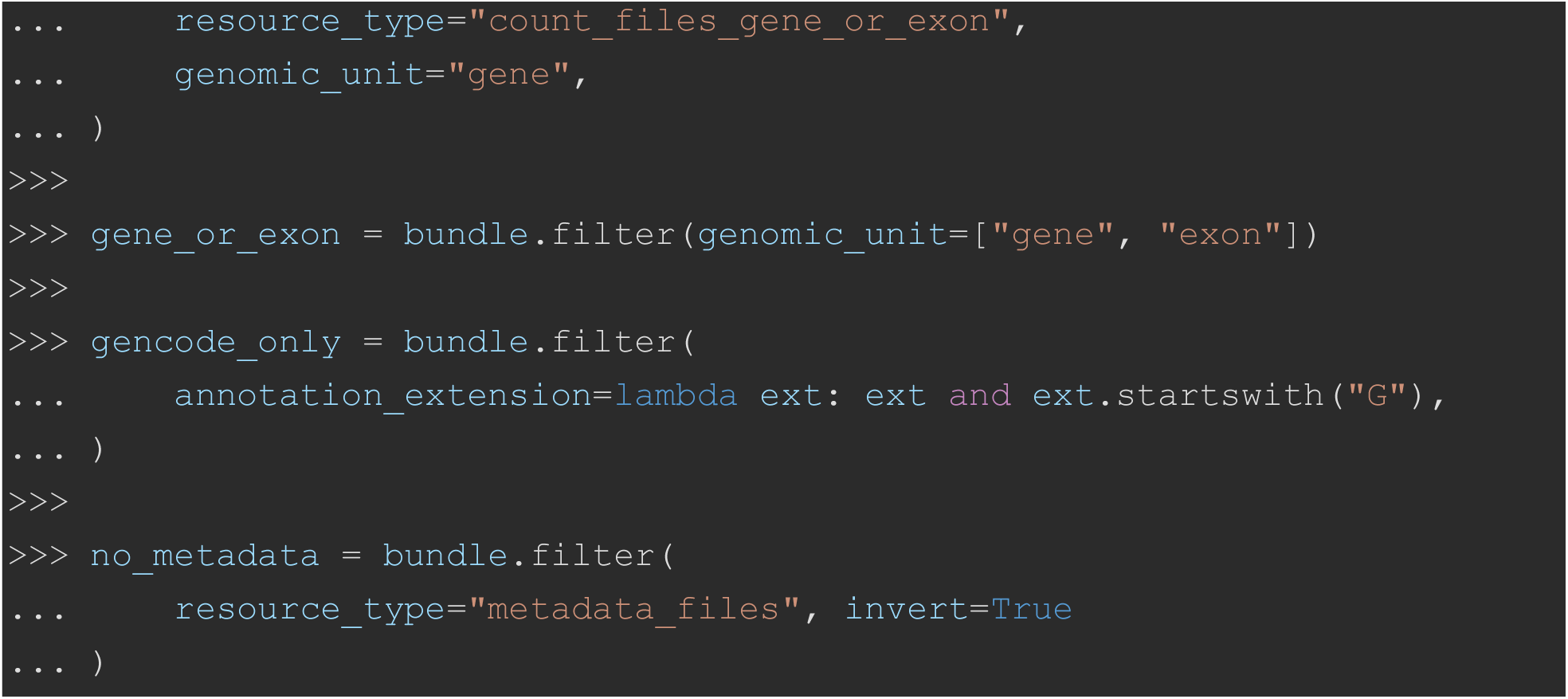

Convenience aliases (only_counts(), only_metadata(), bigwigs(), exclude_metadata()) wrap the most frequent filter combinations:

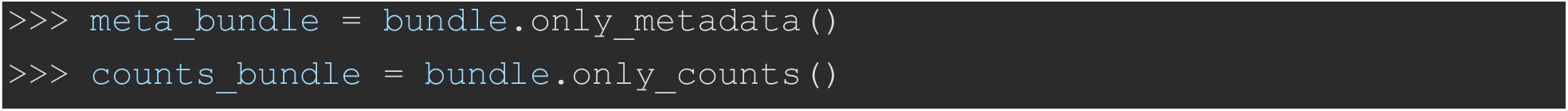

To combine distributed files into a single matrix, the stack_count_matrices() method concatenates compatible DataFrames along a specified axis. Data integrity is maintained via the compat parameter, which prevents the merging of incompatible datasets. Setting this parameter to “feature” ensures that matrices share an identical feature family, the same genomic unit (gene versus exon), or the same junction subtype, before concatenation:

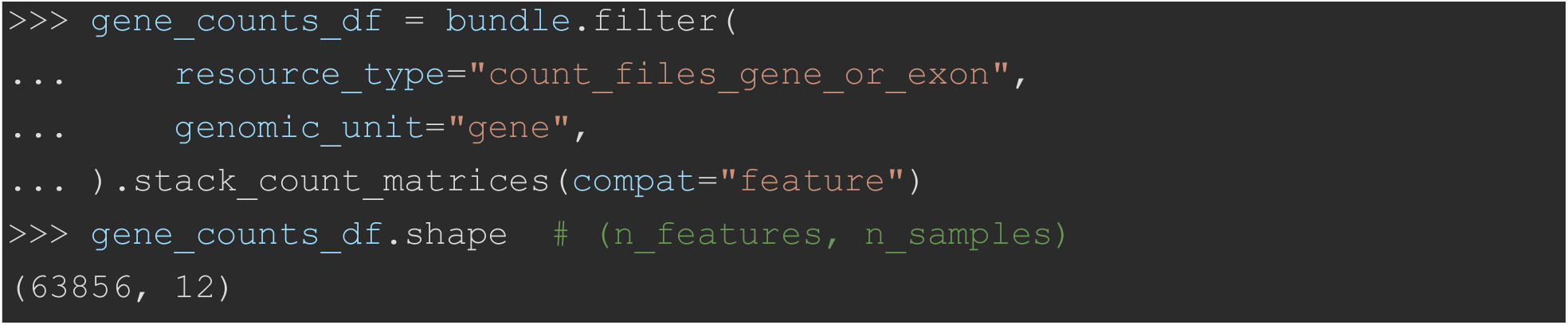

Resource bundles also serve as the input to the BiocPy builder methods to_summarized_experiment() and to_ranged_summarized_experiment(), which the high-level create_rse() function dispatches to internally. Constructing an object directly from a bundle exposes the intermediate stage for users who need to inspect or transform resources between discovery and assembly:

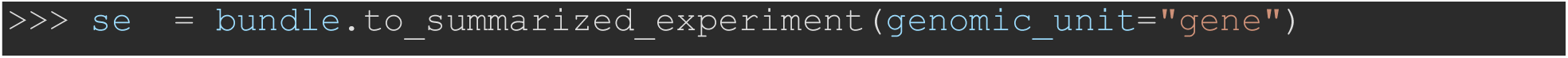
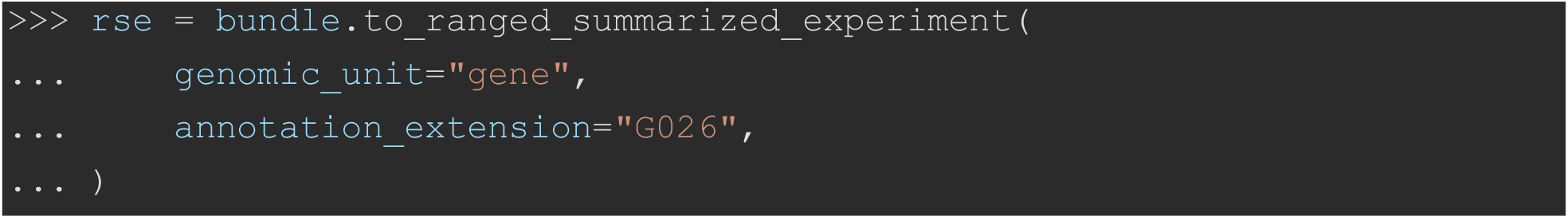

### Metadata processing and count normalization

The recount3 repository distributes base-pair coverage sums rather than absolute read counts. To address this, the recount3.se submodule provides scaling and transformation utilities consistent with the original R implementation. The compute_read_counts() function converts coverage sums into approximate read counts by adjusting for the average mapped read length recorded in the sample metadata:

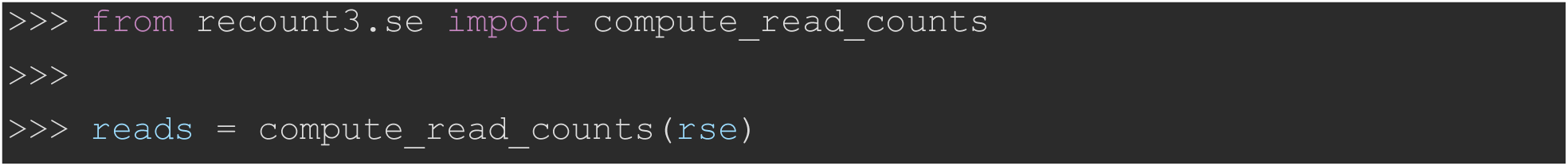

To normalize for library size, compute_scale_factors() calculates per-sample scaling values using either the area under the curve of the coverage (AUC) or mapped-reads method, and transform_counts() applies these factors to the count matrix to produce comparable scaled counts across samples:

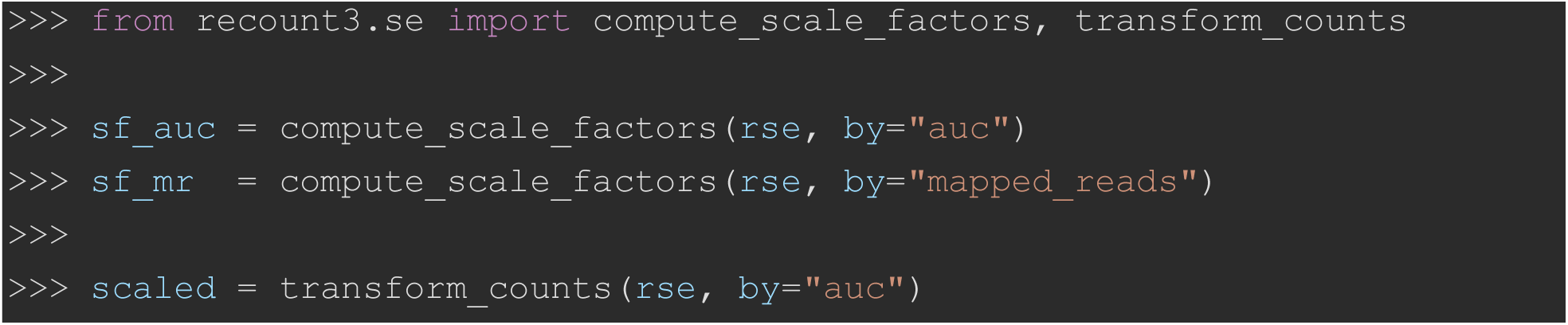

Additionally, compute_tpm() calculates transcripts per million (TPM) using the genomic feature widths attached to the RSE object, and is_paired_end() infers paired-end status from the same metadata columns used by the R-side helper of the same name:

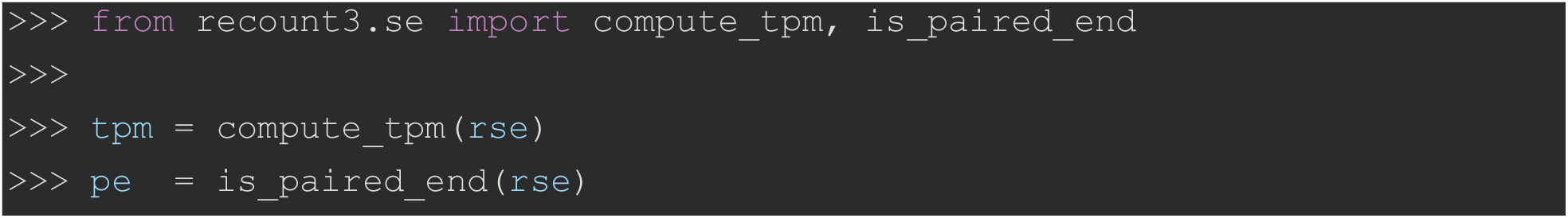

SRA metadata handling is managed by the expand_sra_attributes() function, which parses pipe-delimited SRA attribute strings into distinct tabular columns and appends them to the experimental column data:

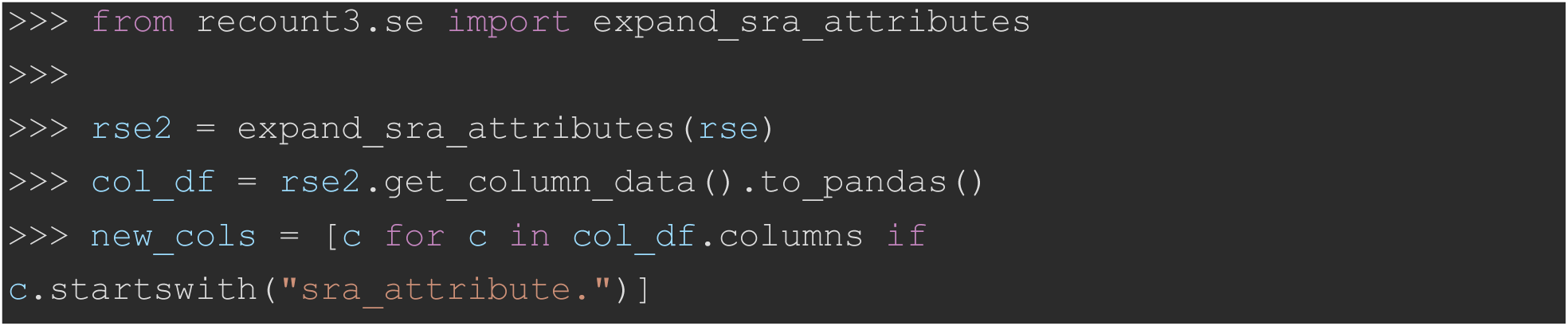

### Low-level resource management and BigWig file access

For tasks requiring direct file access, the R3Resource class controls caching and materialization. Each resource defines a deterministic URL based on the repository’s layout. A description is built via the R3ResourceDescription factory, which routes to the appropriate registered subclass for the requested resource_type:

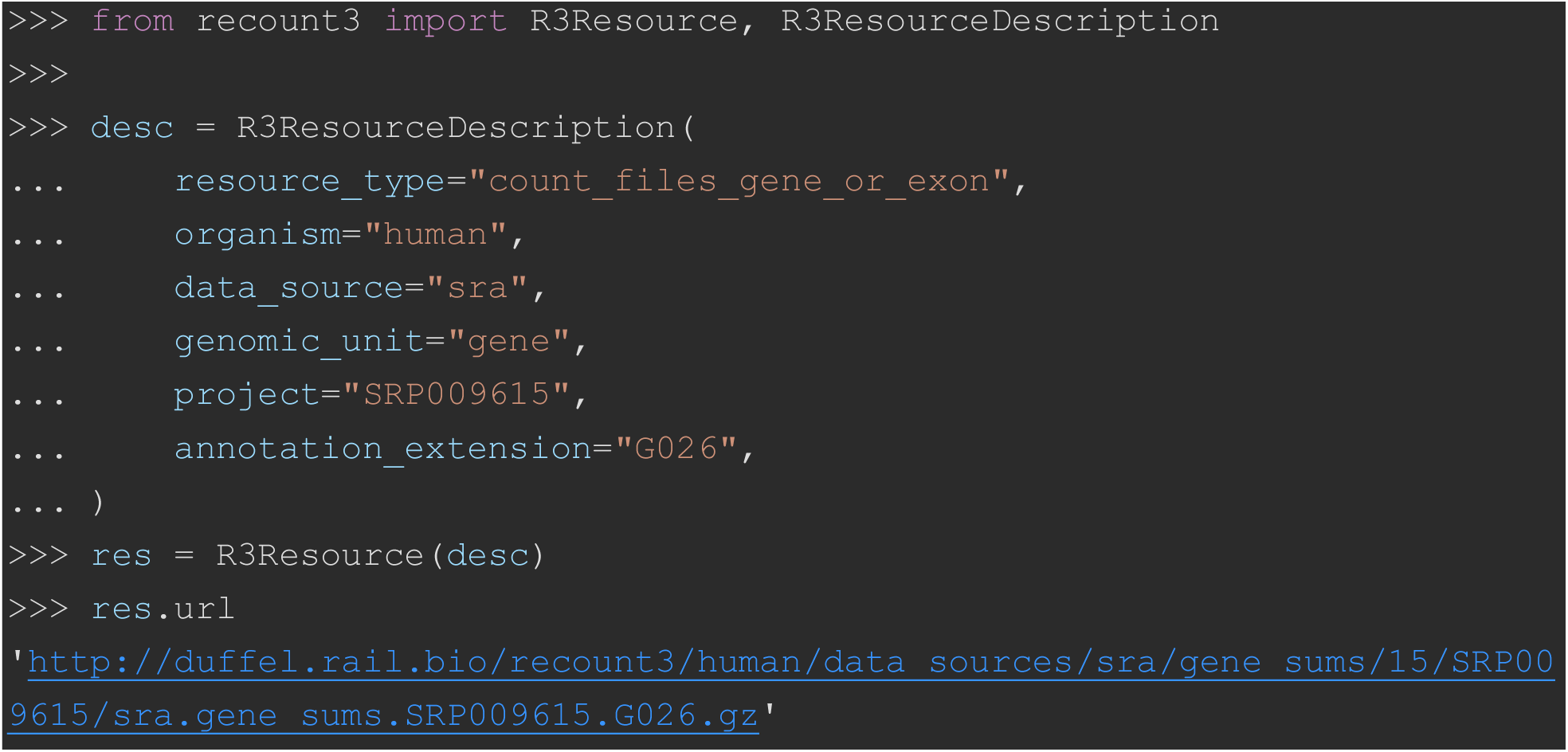

Through the download() method, users specify a local directory path, a .zip archive (entries written with ZIP_DEFLATED compression and stored under their relative URL path), or None for cache-only retrieval. The cache_mode parameter controls cache behavior, allowing users to bypass the cache (“disable”), force a refresh (“update”), or use locally stored files (“enable”, the default):

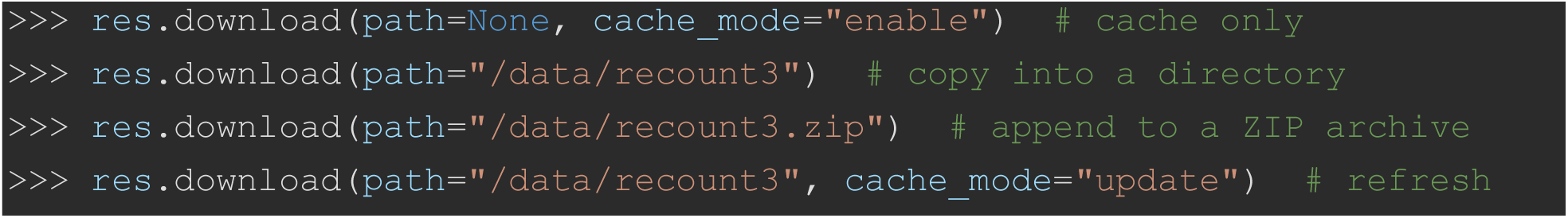

For multi-file retrieval, R3ResourceBundle.download() fetches a bundle’s resources concurrently using a pool of worker threads sized by max_workers (default 8). Because downloading is I/O-bound, this parallelism overlaps the per-file network latency and mirrors, and shares its thread-pool implementation with, the CLI’s --jobs option:

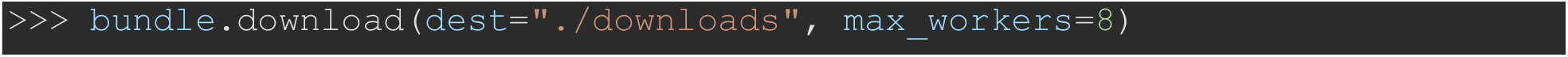

Cache management is exposed through three module-level helpers that operate on the directory pointed to by RECOUNT3_CACHE_DIR (default ∼/.cache/recount3/files). Files in this directory follow a flat {sha256(url)[:16]} {basename} layout, enabling URL-keyed lookups without filesystem hierarchy ambiguity:

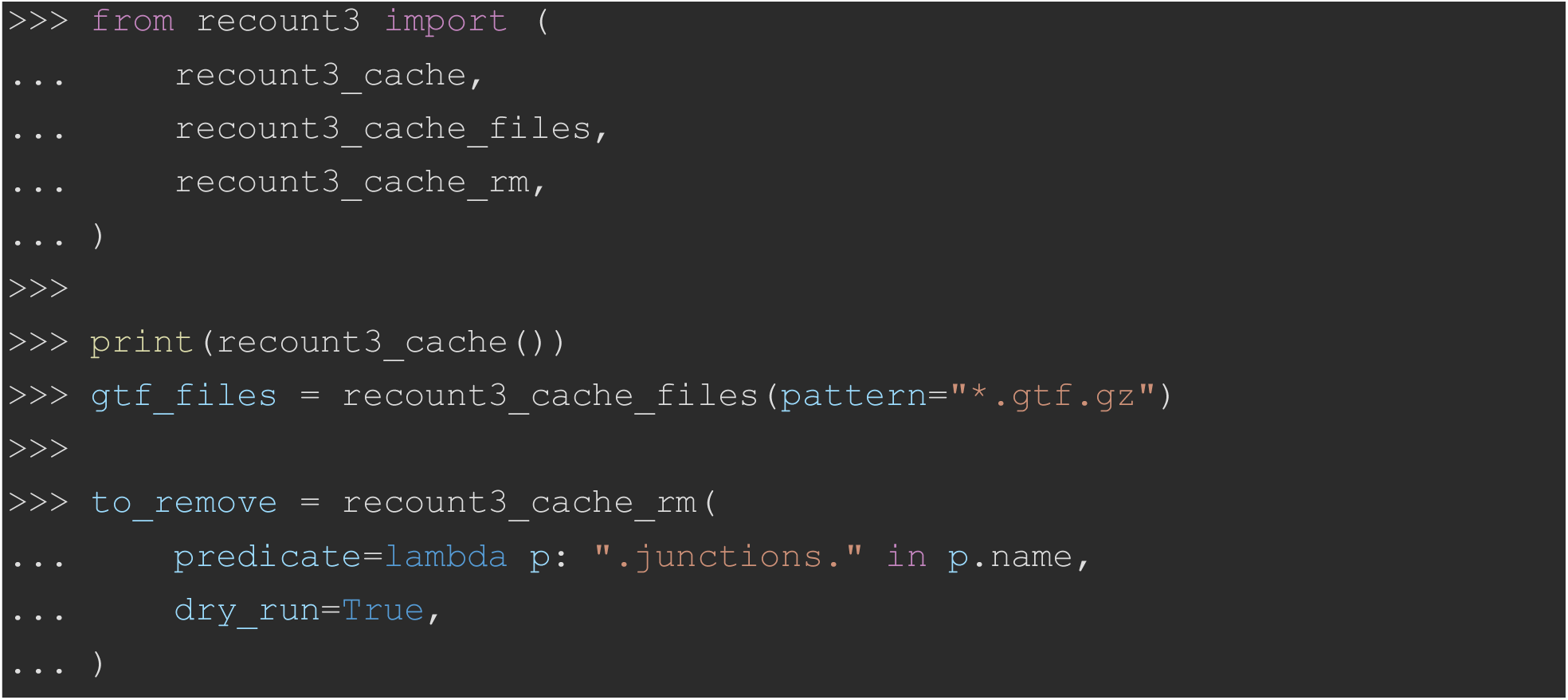

The package also supports the retrieval of per-sample base coverage files. Filtering a bundle for BigWig resources yields file handlers wrapped by the recount3._bigwig.BigWigFile class, which requires the optional pyBigWig dependency [6]. This wrapper lazily loads the file to reduce memory consumption and permits querying of specific chromosomal regions:

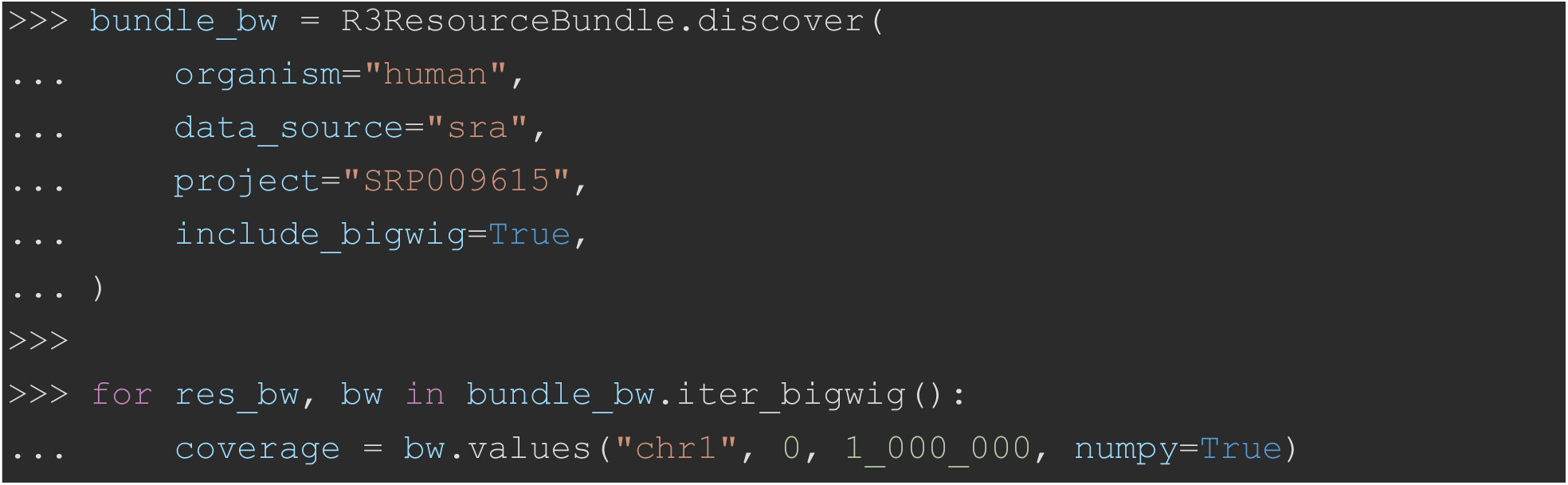
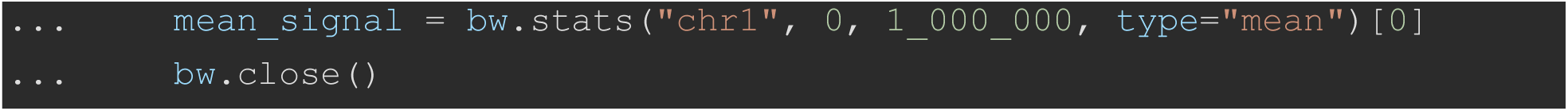

Network and cache behavior is centralized in an immutable Config dataclass, which can be passed to any resource or search function to override the environment-derived defaults read by default_config():

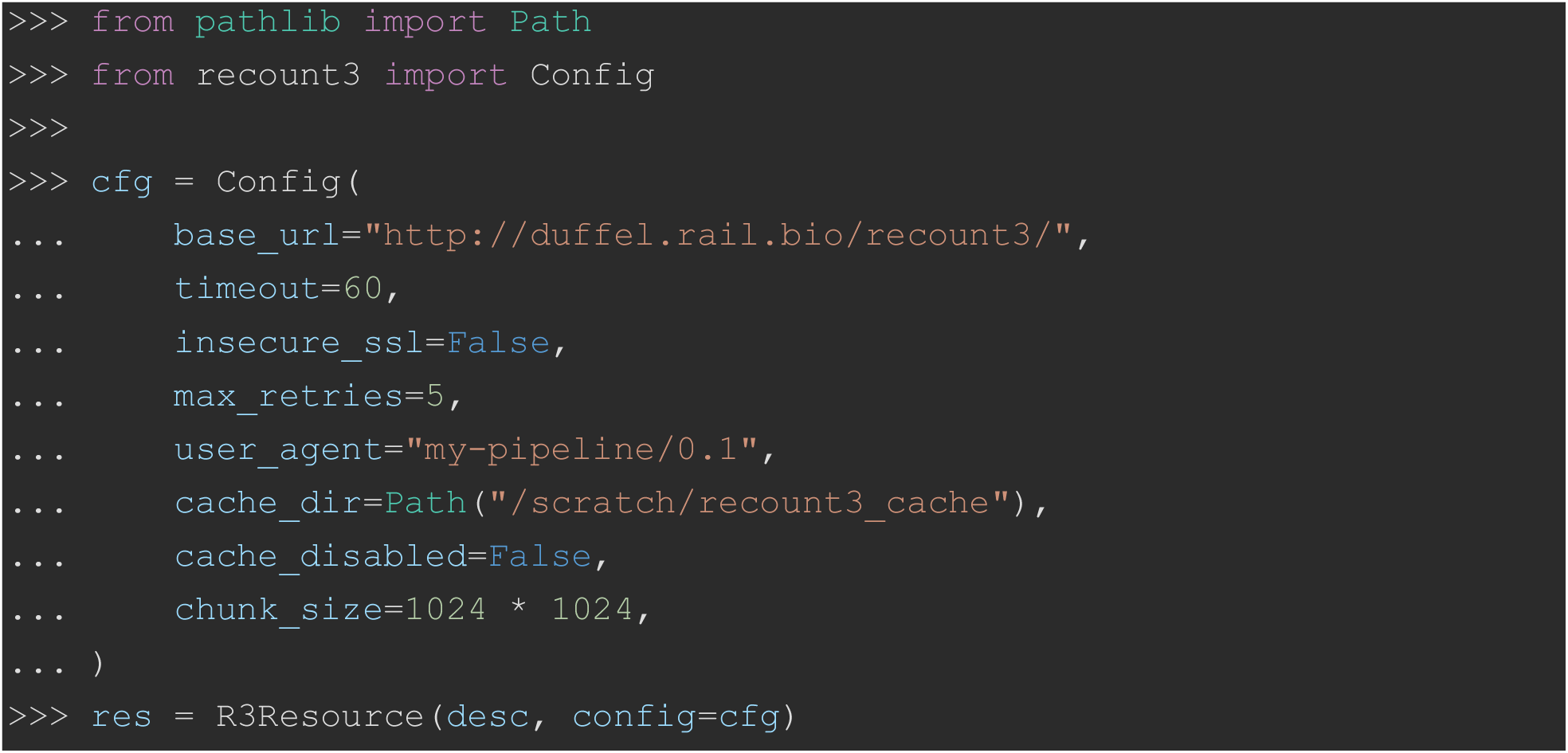

### Command-Line Interface (CLI) for scalable workflows

To support direct utilization on the command line as well as in shell scripts, data analysis pipelines, and complex data workflows that often execute within high-performance computing (HPC) environments, recount3 provides a CLI structured around a discover–manifest–materialize workflow. The search subcommand emits a JSON Lines (JSONL) [7] or tab-separated values (TSV) manifest of required resources, with one structured record per file:

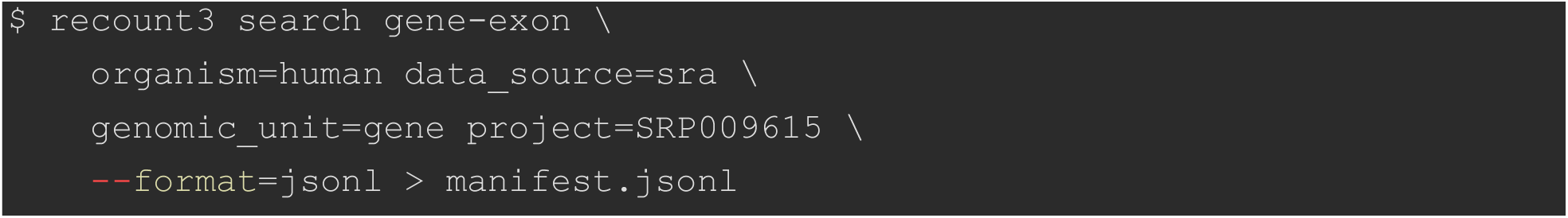

The search modes mirror the API’s resource families (gene-exon, junctions, annotations, metadata, bigwig, sources, source-meta), with a project-wide convenience mode that enumerates every resource for a given study:

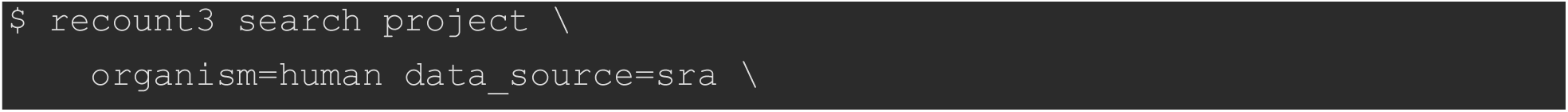
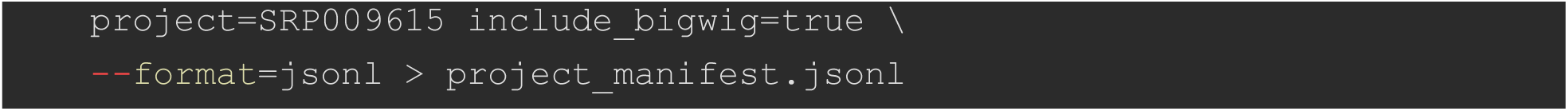

The manifest is passed to the download subcommand, which reads the JSONL stream from a file or standard input and fetches the data using parallel worker threads specified by the --jobs argument:

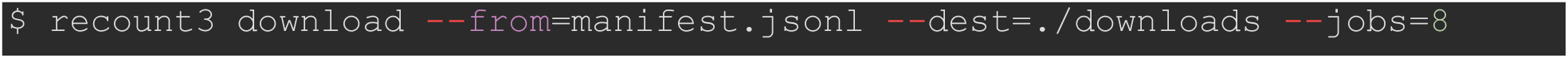

Because both subcommands operate on JSONL via standard streams, search and download can be composed into a single command pipeline without an intermediate file, which is well-suited to streaming workflows:

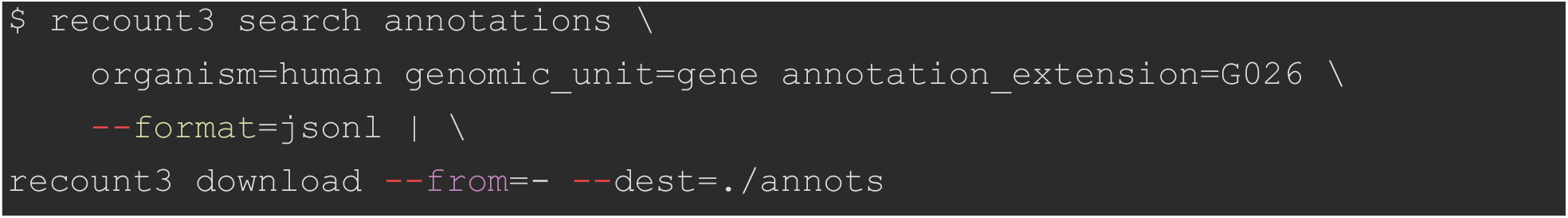

The CLI also supports parallel streaming into a single ZIP_DEFLATED archive and offers cache-mode flags equivalent to the API, allowing pipelines to force fresh downloads when correctness requires bypassing the cache:

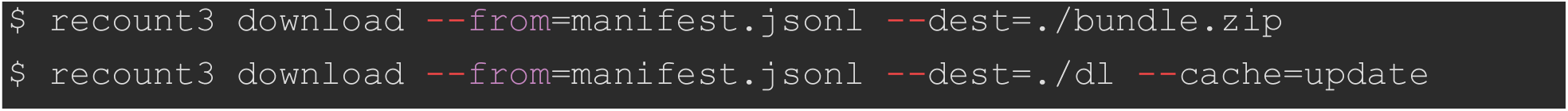

For end-to-end assembly without a Python session, the bundle subcommand group consumes a manifest and produces analysis-ready outputs: bundle stack-counts writes a stacked count matrix as TSV, gzip-compressed TSV, or Apache Parquet [8]; bundle se and bundle rse produce pickled SummarizedExperiment and RangedSummarizedExperiment objects (or AnnData HDF5 files, when the optional anndata dependency is installed):

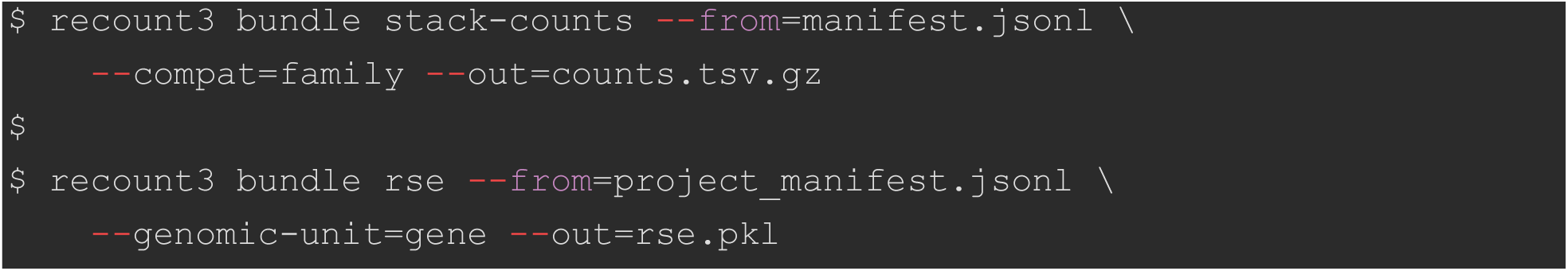

Auxiliary commands round out the CLI surface. The ids subcommand emits unique sample and project identifier lists, which are useful for scripting discovery loops over an entire data source, and the smoke-test subcommand provides a small connectivity probe for continuous-integration and local validation:

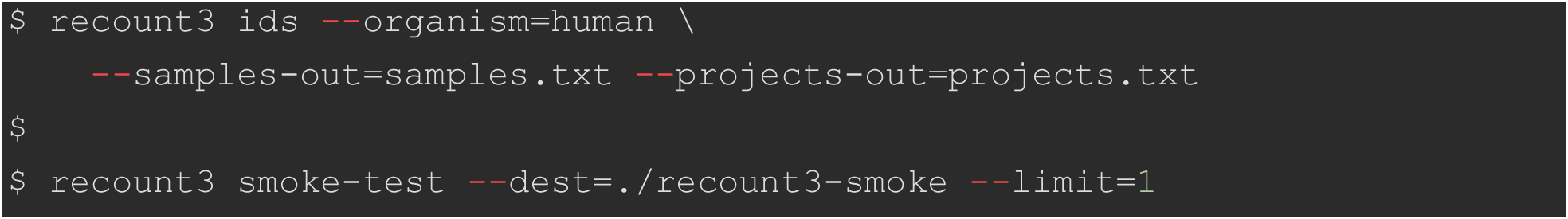

### Parallel download performance

Because file retrieval is I/O-bound, the threaded downloading shared by R3ResourceBundle.download(max_workers=…) and the CLI --jobs option markedly accelerates multi-file retrieval. Benchmarked against the live recount3 mirror, 38 small per-project files (34.9 MB total) at a measured round-trip time of approximately 507 ms, throughput rose from 1.5 MB/s with a single worker to a peak of 6.7 MB/s, a 4.4-fold speedup, with most of the gain already realized by eight workers (Figure 5). The curve flattens beyond this knee, reflecting per-file connection overhead and, for larger transfers, saturation of the network link; the default of eight workers therefore captures the bulk of the attainable speedup under typical download use-case conditions.

Throughput is the total bytes transferred divided by wall-clock time; markers show the mean over four replicates and error bars the estimated 95% confidence interval. Throughput increases steeply through approximately eight threads and then plateaus.

## Discussion

### Implications for large-scale RNA-seq reuse

The principal purpose of this work is to place recount3’s curated corpus within immediate reach of the Python scientific stack. The package returns standard numpy [9], pandas [10], and scipy [11] objects, with retrieved count matrices and metadata flow without conversion into the downstream Python ecosystem for statistical analysis, visualization, machine learning, and deep learning, including single-cell-oriented tooling such as scanpy and the AnnData format [12]. The three-tier API is designed so that this reach does not come at the cost of accessibility: create_rse offers one-call assembly of an analysis-ready object for newcomers, R3ResourceBundle supports multi-project discovery and curated subsetting for power users, and R3Resource exposes per-file caching and materialization for workflows that require fine-grained control. This layering allows a researcher to begin with a single high-level call and progressively adopt lower-level constructs only as required. The CLI broadens access further still. The CLI’s discover–manifest–materialize model brings recount3 to non-Python users and to batch and HPC environments where a Python session is inconvenient, and the JSON Lines (JSONL) manifests serve a dual role as shareable, reproducible provenance artifacts that decouple the act of discovery from the act of download.

### Interoperability, Reproducibility, and Maintainability

By constructing BiocPy SummarizedExperiment, RangedSummarizedExperiment, and GenomicRanges objects [13], the package mirrors the central data structures of Bioconductor, easing migration of established Bioconductor workflows into Python and supporting interoperability and reproducibility across the two languages; optional AnnData export extends this reach toward the single-cell and scanpy ecosystems. This cross-language fidelity is reinforced by a faithful re-implementation of recount3’s R-side semantics, including coverage-sum scaling by the area under the curve of coverage (AUC) or mapped reads, transcripts-per-million (TPM) computation, paired-end inference, SRA attribute expansion, and the raw_counts/counts assay-naming convention. Therefore, mixed R and Python teams can expect numerically consistent results (summarized in the Python-versus-R feature comparison of Table 1). Several engineering choices further support reuse and long-term maintainability: comprehensive PEP 561 type annotations propagate type information to downstream code, lazy optional-dependency imports surface actionable installation guidance only when a feature is actually used, deterministic URL construction makes resource identity reproducible, and a persistent on-disk cache avoids redundant transfers. Within the landscape of Python access tools, the package occupies a distinct niche: it is not a low-level coverage reader such as pyBigWig [6] (which it wraps), nor a generic repository-metadata client such as GEOparse [14], nor an alternative uniformly-processed expression portal such as ARCHS4 [15]; rather, it provides recount3-specific, manifest-driven discovery integrated with the BiocPy object model.

**Table 1.** Feature comparison of the recount3 Python package to the recount3 R package.

| **Feature** | **recount3 (Python)** | **recount3 (R)** |
| --- | --- | --- |
| Programming language | Python | R |
| Distribution channel | PyPI (pip install recount3) | Bioconductor<br>(BiocManager::install) |
| Type annotations | Yes (PEP 561) | No |
| Command-line interface | Yes | No |
| Discover resources for a project | R3ResourceBundle.discover() | available_projects() /<br>available_samples() |
| Multi-project bundling | Yes (project=[...] Cartesian product) | Partial (loop / lapply by user) |
| Parallel downloads | Yes (API max_workers, CLI --jobs; threaded) | No |
| ZIP archive utilization | Yes | No |
| On-disk file cache | Yes | Yes |
| Manifest output | Yes (JSONL and TSV) | No |
| RangedSummarizedExperiment from project | create_rse() | create_rse() |
| SummarizedExperiment from custom inputs | R3ResourceBundle.to_summarized_experiment() | create_rse_manual() |
| Gene/exon counts | Yes | Yes |
| Junction counts (MM/ID/RR) | Yes | Yes |
| BigWig coverage | Yes (pyBigWig) | Yes (rtracklayer) |
| GTF annotation parsing | Yes | Yes |
| SRA attribute expansion | `expand_sra_attributes()` | `expand_sra_attributes()` |
| Paired-end inference | `is_paired_end()` | `is_paired_end()` |
| TPM computation | `compute_tpm()` | `recount::getTPM()` |
| AUC-based scaling | `compute_scale_factors() /`<br>`transform_counts()` | `compute_scale_factors() /`<br>`compute_scaled_counts()` |
| Mapped-reads scaling | `compute_scale_factors(by="`<br>`mapped_reads")` | `compute_scale_factors(by="`<br>`mapped_reads")` |

### Performance and operational considerations

Because file retrieval from the recount3 mirror is input/output-bound, the package overlaps per-file network latency using a thread pool, which is effective because CPython releases the global interpreter lock during socket and disk input/output. The resulting throughput gains are quantified in the Parallel download benchmark, where most of the attainable speedup is realized by roughly eight workers utilizing a 100Mbit/s internet connection. Therefore, the shared default of eight workers for both R3ResourceBundle.download() and the command-line --jobs option is a deliberately conservative operating point. In practice, we recommend keeping the on-disk cache enabled and bounding concurrency for routine work, both to accelerate repeated workflows and as a matter of good citizenship toward a shared public mirror. Also, following good internet citizenship should prevent the necessity of blacklisting. Next, materialization hard-links cached files into a destination when the filesystem permits, avoiding a redundant second write, and the cache-management helpers together with the selectable cache modes allow users to force fresh downloads when correctness requires bypassing the cache. Two operational caveats deserve emphasis for prospective users. First, the interoperability and coverage features depend on optional packages, the BiocPy stack for SummarizedExperiment construction and pyBigWig for coverage access, whose compiled components are less portable to non-Linux platforms; the package isolates these as optional dependencies and degrades gracefully in their absence. The test suite runs in continuous integration on Linux across Python 3.10 through 3.14 and on macOS and Windows at the endpoint versions, 3.10 and 3.14. On macOS the BiocPy stack is exercised but pyBigWig is omitted, while on Windows, where neither pyBigWig nor the BiocPy ranged-coordinate dependency provides a binary wheel, the suite installs the pure-Python core and automatically skips the tests that require those native extensions. Second, all network and cache behavior is centralized in a single immutable configuration object and mirrored by environment variables and command-line flags, so deployment in constrained environments (custom mirrors, proxies, or strict TLS policies) is configurable without code changes. A practical obstacle during development was that recount3’s own documentation of its file-layout and URL conventions is sparse and, in places, incorrect. For example, the published annotation-code table transposes two build identifiers, and the documented per-sample coverage-path rule does not hold for GTEx, whose shard is drawn from an interior offset of the sample identifier rather than its final characters. The package’s URL templates were therefore established empirically, cross-checked against the recount3 R reference implementation and validated for every resource type against the live mirror.

### Limitations

Several limitations bound the scope of this contribution and should inform its use. Most fundamentally, the package is an access and assembly layer, not a sequence-processing pipeline: it retrieves and organizes the outputs of recount3’s uniform processing but does not realign or reprocess raw reads, and it therefore inherits recount3’s upstream decisions regarding alignment, annotation, and quality control. Its coverage is likewise bounded by what the recount3 online resource provides, human and mouse only, across the SRA, GTEx, and TCGA data sources, and limited to the annotation builds that recount3 distributes. The package is also coupled to recount3’s file-layout convention through deterministic URL templates, so a future reorganization of that layout would break discovery and require updated resource descriptors. The extensible registry-based factory keeps the addition of new resource types localized should the layout evolve.

Further limitations concern the derived quantities and the functional surface. Genomic-range resolution depends on concordance between annotation feature identifiers and count-matrix identifiers, reconciled where possible by an Ensembl version-stripping fallback, and, for junctions, on row agreement between the matrix and its row-ranges sidecar; when ranges cannot be resolved, the package either falls back to a plain SummarizedExperiment or raises an informative error. In terms of functional scope, the package deliberately delegates differential expression analysis to downstream Python and Bioconductor tools rather than providing it directly, and it does not replicate snapcount-style fast querying of arbitrary genomic intervals across the corpus. Finally, the performance characterization, consistent with the Methods and Results, rests on a small, deliberately bounded workload chosen to avoid burdening the public mirror, a single real-network condition, and throughput that is not portable across networks and may be shaped by server-side connection limits.

### Comparison to R package

As shown in Table 1, the Python recount3 package provides all of the functionality in the R recount3 package. Moreover, the Python recount3 package provides a variety of new functionalities. The most important new functionality is the CLI, which is very useful for enabling shell scripting as well as facilitating utilization of the recount3 online resource within data analysis pipelines and workflow managers. The next functionality is the use of multithreading and parallelized downloads that greatly improve download throughput. The Python recount3 package also allows use of zip archives for resource storage and manifestation. Also, the package can manifest output in JSON Lines and TSV formats. Lastly, the package provides Python typing annotations (PEP 561), which helps improve the quality of derivative code that imports and uses the package.

### Future directions

Several extensions follow naturally from the present design. As the recount3 resource evolves, through additional data sources or organisms, or a future major release, the registry-based factory (see Methods) makes supporting new resource types a low-cost, localized change that does not disturb existing code. Deeper integration with the AnnData and scanpy ecosystems, along with machine-learning-ready export paths, would further lower the barrier between the recount3 corpus and modern analysis frameworks. The manifest-centric command-line design is also a natural fit for workflow managers such as Snakemake [16] or Nextflow [17], and could be complemented in future by snapcount-style remote interval querying, lazy or remote slicing of BigWig coverage, and cloud or object-store cache backends for large shared deployments. Support for collections, manually curated set of samples spanning one or more studies that includes collection-specific sample metadata, is anticipated once collections become available on the “Duffel” mirror, whereas they currently only exist on a separate Snaptron server (http://snaptron.cs.jhu.edu/data/temp/recount3), which is not prominently labeled as a recount3 mirror for public use. The package’s comprehensive typing and modular architecture are intended to keep the barrier to community contribution low as these directions are pursued.

### Conclusions

The recount3 Python package delivers the first general-purpose Python API and CLI for the recount3 resource, complementing rather than replacing the established recount3 and snapcount R/Bioconductor packages. By exposing the same uniformly-processed human and mouse RNA-seq corpus through idiomatic Python objects and a scriptable command line, the package extends large-scale data reuse beyond the R/Bioconductor user base to the much larger Python data-science and machine-learning community, and to language-agnostic shell and high-performance computing (HPC) pipelines. In doing so, it directly addresses the reuse barrier emphasized in the Introduction: the practical difficulty of programmatically discovering, retrieving, and assembling tens of thousands of pre-processed datasets without bespoke tooling.

## Methods

### Implementation and architecture

The recount3 package is implemented in the Python programming language (version 3.10 or higher) with a modular, object-oriented architecture. The public surface is organized into focused submodules (see Figure 1). The recount3.config submodule centralizes runtime configuration as an immutable Config dataclass. The recount3.resource submodule defines R3Resource, the class that owns each data file’s lifecycle. The recount3.search submodule provides query and discovery functions. The recount3.bundle and recount3.se submodules provide aggregation facilities that compose single-file resources into analysis-ready objects. The recount3.errors submodule defines a small hierarchy of domain-specific exceptions rooted in Recount3Error, and the recount3.types submodule exposes the type aliases (CacheMode, CompatibilityMode, FieldSpec, StringOrIterable) that parameterize the public API. The recount3.cli submodule implements the command-line entry point (see “Command-line interface” below). Three private submodules support the public ones: recount3._descriptions defines resource description dataclasses and uniform resource locator (URL) resolution, recount3._utils provides file-system access, compression, caching, and HTTP utilities, and recount3._bigwig wraps the optional pyBigWig library. The package is fully typed and distributes a PEP 561 py.typed marker so that downstream code receives the same type hints used internally.

**Figure 1.**
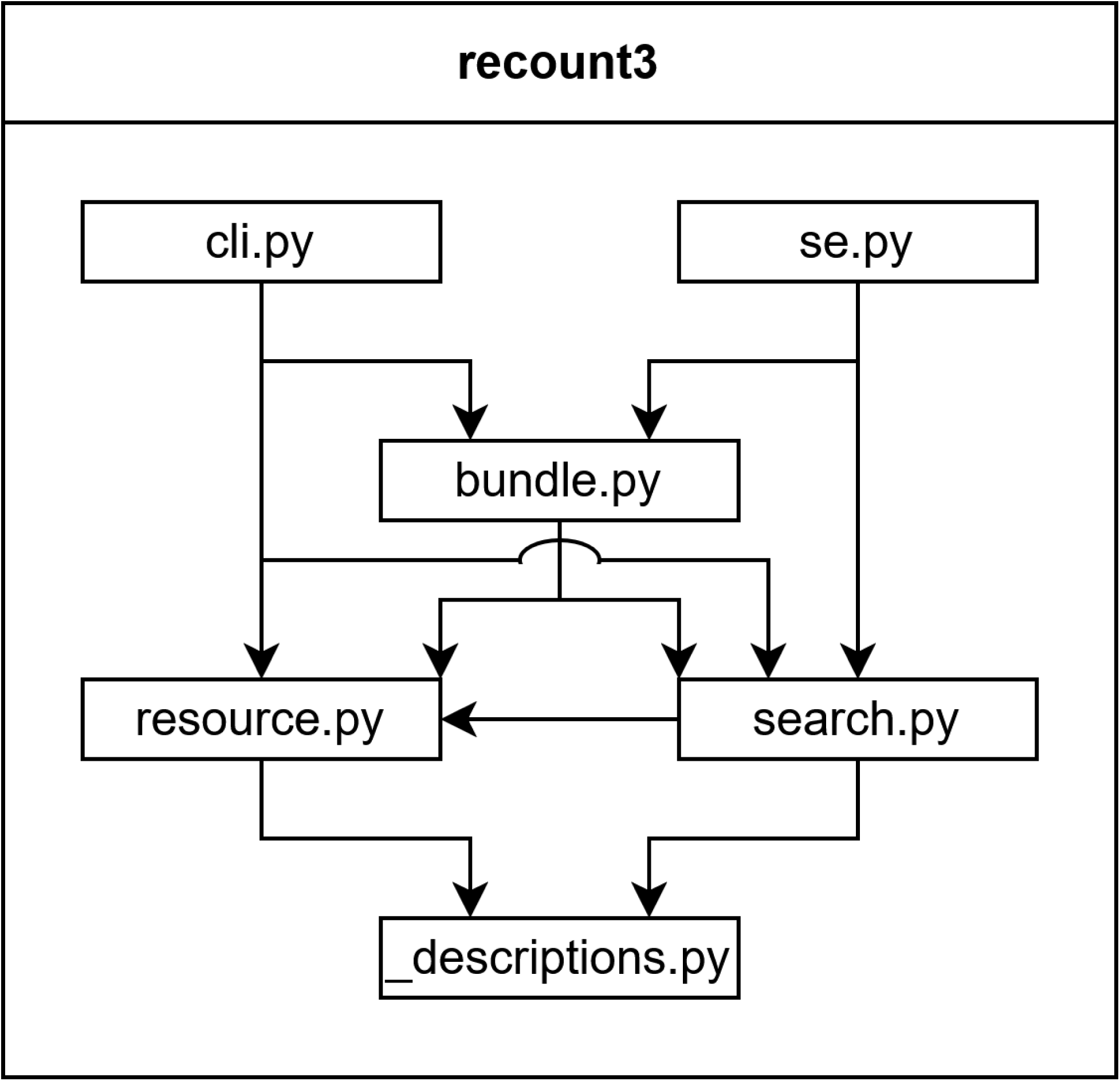
UML package diagram with submodule dependencies. The _bigwig.py and _utils.py private submodules are left out for simplification to major dependencies.

The core object-oriented unit of the package is the R3Resource class, defined in the recount3.resource submodule. Each R3Resource instance manages the lifecycle of a single recount3 data file, encompassing URL resolution, network retrieval, local caching, and materialization (loading, parsing, and transformation of a recount3 data file into an in-memory object). The R3ResourceBundle class, defined in the recount3.bundle submodule, is a container class. Each R3ResourceBundle instance object manages a collection of R3Resource instance objects and exposes filtering, lazy loading, and BiocPy construction helpers. Figure 2 shows the inheritance and aggregation relationships between these classes.

**Figure 2.**
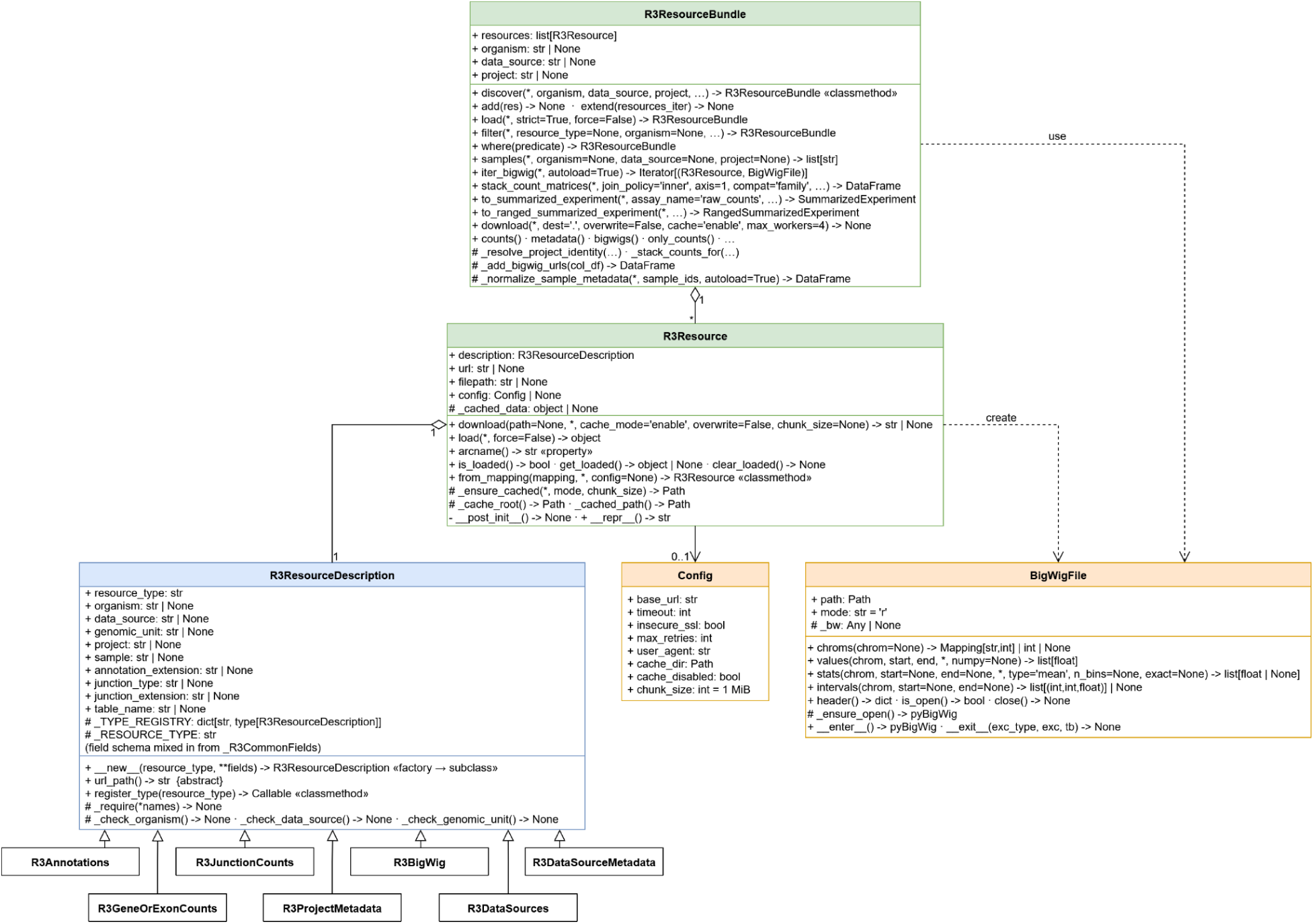
UML class diagram with class dependencies.

### Resource description and URL resolution

The recount3 repository provides structured access of data files organized by study (referred to as a project) that is similar to a REST interface [18]. This access requires deterministic URL construction matching recount3’s published file layout. The same relative layout is served by each of recount3’s public mirrors; by default the package targets the “Duffel” load-balancer host (duffel.rail.bio), but any mirror sharing this layout, for example the AWS Open Data registry or the JHU IDIES Dataverse, can be selected through the configurable base URL alone (see “Downloading, caching, and materialization”).

The private recount3._descriptions submodule defines a series of dataclasses [19] that inherit from the abstract R3ResourceDescription class and a shared _R3CommonFields mixin class. Each concrete dataclass acts as a resource descriptor that abstracts the URL schema and the URL-construction task, allowing users to request data by intuitive biological concepts (e.g., organism=“human”, project=“SRP009615”) rather than requiring the manual construction of exact file-path-like URLs.

R3ResourceDescription embodies a registry-based factory design pattern, fusing aspects of a registry design pattern [20] with the Gang of Four abstract factory design pattern [21], which produces a concrete product without requiring client code to know which subclass to instantiate. Two design choices distinguish it. First, the abstract base R3ResourceDescription is itself the dispatcher: a single R3ResourceDescription(resource_type=…) call returns an instance of the matching subclass, rather than requiring the client to first select a concrete factory and then call product-creation methods on it. Second, concrete subclasses are not hard-coded into the dispatcher. Each subclass registers itself at import time via the @R3ResourceDescription.register_type(“resource_type_tag”) class decorator, which inserts an entry into a class-level mapping _TYPE_REGISTRY: dict[str, type[R3ResourceDescription]] and stamps the subclass with its registration key. When the abstract base is instantiated, its custom new looks up the requested resource_type in _TYPE_REGISTRY and routes construction to the matching subclass; instantiating a concrete subclass directly bypasses the registry lookup. This design is open for extension: new resource types can be added without modifying the dispatcher, making _TYPE_REGISTRY.keys() the authoritative list of supported resource types.

The shared field surface is supplied by _R3CommonFields, a mixin, slots-enabled dataclass that declares every parameter any descriptor might need (resource_type, organism, data_source, genomic_unit, project, sample, annotation_extension, junction_type, junction_extension, table_name). Concrete subclasses inherit from both _R3CommonFields and R3ResourceDescription, so generic code (serialization, filtering, manifest emission) can read the same attribute names from any descriptor without special-casing the resource type. Although every field is present on every descriptor, only a subset is required for a given resource type; each subclass’s post_init calls a _require(…) helper to enforce its own required fields and invokes type-specific validators that reject invalid organisms, data sources, or genomic units. Misconfigured parameters therefore surface at object construction time, before any HTTP request is initiated.

Supported file types managed by these resource descriptors include gene and exon count matrices in tabular files, junction counts in MatrixMarket format [22], project metadata in tabular files, organism-level data-source indices, and sample-level BigWig coverage files. Gzip-compressed tabular files are handled transparently. Annotation gene transfer format (GTF) files are also fetched through the same description and caching machinery, but their parsing is performed internally by the bundle layer during RangedSummarizedExperiment construction rather than through the public R3Resource.load() method. Table 2 lists the resource types and their description subclasses, with Supplemental Table 1 providing the URL template for each resource type.

**Table 2.** The recount3 resource types and their associated R3ResourceDescription subclasses.

| Resource Type | Description Class | File Format | In-Memory Representation | Required Fields | Granularity |
| --- | --- | --- | --- | --- | --- |
| annotations | R3Annotations | GTF (gzip) | pandas.DataFrame | organism,<br>genomic_unit,<br>annotation_extension | organism<br>+<br>annotation |
| count_files_gene_or_exon | R3GeneOrExonCounts | TSV (gzip) | pandas.DataFrame | organism, data_source, genomic_unit, project, annotation_extension | project |
| count_files_junctions (MM) | R3JunctionCounts | MatrixMarket (gzip) | pandas.DataFrame (sparse) | organism, data_source, project, junction_type, junction_extension=MM | project |
| count_files_junctions (ID) | R3JunctionCounts | TSV (gzip) | pandas.DataFrame | organism, data_source, project, junction_type, junction_extension=ID | project |
| count_files_junctions (RR) | R3JunctionCounts | TSV (gzip) | pandas.DataFrame | organism, data_source, project, junction_type, junction_extension=RR | project |
| metadata_files | R3ProjectMetadata | TSV (gzip) | pandas.DataFrame | organism, data_source, project, table_name | project x table |
| bigwig_files | R3BigWig | BigWig (.bw) | recount3._bigwig.BigWigFile | organism, data_source, project, sample | sample |
| data_source_metadata | R3DataSourceMetadata | TSV (gzip) | pandas.DataFrame | organism, data_source | organism + data source |
| data_sources | R3DataSources | plain text | none | organism | organism |

### Search and discovery

The recount3.search submodule provides functions to query available projects (i.e., studies), enumerate samples, and resolve annotation identifiers without requiring the download of large data matrices. Tier-one search functions (search_count_files_gene_or_exon, search_count_files_junctions, search_metadata_files, search_bigwig_files, search_annotations, search_data_sources, search_data_source_metadata, and the omnibus search_project_all) return collections of R3Resource objects. Tier-two helpers (available_samples, available_projects, project_homes, samples_for_project, annotation_options, and annotation_ext) load and parse the small index files needed to translate human-readable queries into the structured parameters consumed by the resource layer.

To streamline high-throughput analysis, every tier-one search function accepts its parameters either as a single string or as an iterable of strings, and computes the Cartesian product of all parameter values internally. For instance, providing a list of three project IDs and two annotation extensions yields the six corresponding R3Resource objects, one for each combination. This eliminates the need to write nested loops in user scripts to fetch multiple datasets.

For advanced workflows, the R3ResourceBundle class acts as a container for grouped resources. R3ResourceBundle instance objects support lazy loading and metadata-based subsetting using the filter() and where() methods. The filter() method accepts a uniform specification for every description field (i.e., a FieldSpec) in the form of a string, an iterable of strings, or a callable predicate, and additionally accepts a free-form predicate callable for selection logic that does not map onto a single field. Convenience aliases (only_counts, only_metadata, bigwigs, exclude_metadata) wrap the most frequent filter combinations, enabling precise control over the data prior to materialization.

### Downloading, caching, and materialization

Data retrieval is executed by the download() method of the R3Resource class, which fetches remote files via HTTP and stores them in a persistent, on-disk cache. The default cache directory is ∼/.cache/recount3/files, and is configurable through the Config dataclass in recount3.config or via environment variables (RECOUNT3_CACHE_DIR, RECOUNT3_URL, RECOUNT3_HTTP_TIMEOUT, RECOUNT3_MAX_RETRIES, RECOUNT3_INSECURE_SSL, RECOUNT3_USER_AGENT, RECOUNT3_CHUNK_SIZE, RECOUNT3_CACHE_DISABLE). Cache entries follow a flat naming scheme, each file is stored as {sha256(url)[:16]} {basename} in a single directory, so that lookups are URL-keyed and free of filesystem-hierarchy ambiguity.

The download path of R3Resource accepts a directory destination, a .zip archive destination, or None (cache-only). The cache_mode parameter selects between using the cache (“enable”), forcing a fresh download (“update”), and bypassing the cache entirely (“disable”). Cache operations are coordinated by a module-level threading.Lock object so that concurrent downloads cannot corrupt a shared cache entry. ZIP-archive materialization is coordinated by per-path locks held in a weakref.WeakValueDictionary object, allowing many resources to stream into the same archive in parallel without inter-archive contention while still preventing intra-archive corruption. Archives are written with zipfile.ZIP_DEFLATED compression. When materializing data to a directory, the package hard-links the cached file into the destination when the filesystem permits and falls back to copying otherwise, avoiding the cost of a second write whenever possible.

The R3ResourceBundle.download() method retrieves all of a bundle’s member resources concurrently through a concurrent.futures.ThreadPoolExecutor object sized by its max_workers parameter (default 8). Because file retrieval is I/O-bound, the interpreter releases the global interpreter lock while blocked on network and disk I/O, this thread-based concurrency overlaps the per-file network latency and yields substantial speedups for multi-file downloads. The command-line download subcommand exposes the identical mechanism through its --jobs flag (also defaulting to 8), so the API and CLI share one implementation strategy; the “Parallel download benchmark” subsection in the Results quantifies the improved throughput.

### Data parsing, aggregation, and BiocPy integration

Upon retrieval, the load() method parses cached files into the appropriate Python objects. Gene and exon count tables are loaded as pandas.DataFrame objects; MatrixMarket-format junction counts are loaded as sparse-backed pandas.DataFrame objects (constructed from a SciPy CSR matrix via pandas.DataFrame.sparse.from_spmatrix); junction sidecar tables (ID, RR) and metadata tables are loaded as pandas.DataFrame objects; and BigWig files are wrapped by recount3._bigwig.BigWigFile objects, which lazily opens the underlying pyBigWig package [6] handle on first use.

The recount3.bundle and recount3.se submodules provide methods to concatenate single-resource objects into unified dataset objects. The stack_count_matrices() method concatenates count data across multiple samples or projects, enforcing two configurable compatibility modes. The compat=“family” mode requires that all inputs belong to the same high-level family (gene/exon versus junctions), rejecting, for example, gene counts mixed with junction counts.

The stricter compat=“feature” mode additionally requires an identical feature family, distinguishing gene-level from exon-level matrices (and, for junctions, the junction subtype), before concatenation. A common challenge in RNA-seq analysis is the manual alignment of sample metadata, count matrices, and genomic coordinates. The create_rse() function addresses this with a single orchestrated call: given a project identifier, organism, and annotation label (or extension code), it internally constructs an R3ResourceBundle, stacks the relevant matrices, merges and namespaces the five per-project metadata tables onto the count columns, and resolves genomic ranges from a standard annotation file (GTF for gene and exon assemblies; the row-ranges “RR” sidecar for junctions). The result is a fully annotated RangedSummarizedExperiment object ready for differential expression analysis. The default junction_extensions are unit-aware so that the junction case includes the RR sidecar automatically. For workflows that need to inspect or transform a bundle between discovery and assembly, the same logic is exposed at the bundle layer via R3ResourceBundle.to_summarized_experiment and R3ResourceBundle.to_ranged_summarized_experiment, and as the bundle-taking wrappers build_summarized_experiment(bundle, …) and build_ranged_summarized_experiment(bundle, …) in recount3.se. Figure 3 illustrates the steps performed during the creation of a RangedSummarizedExperiment object.

**Figure 3.**
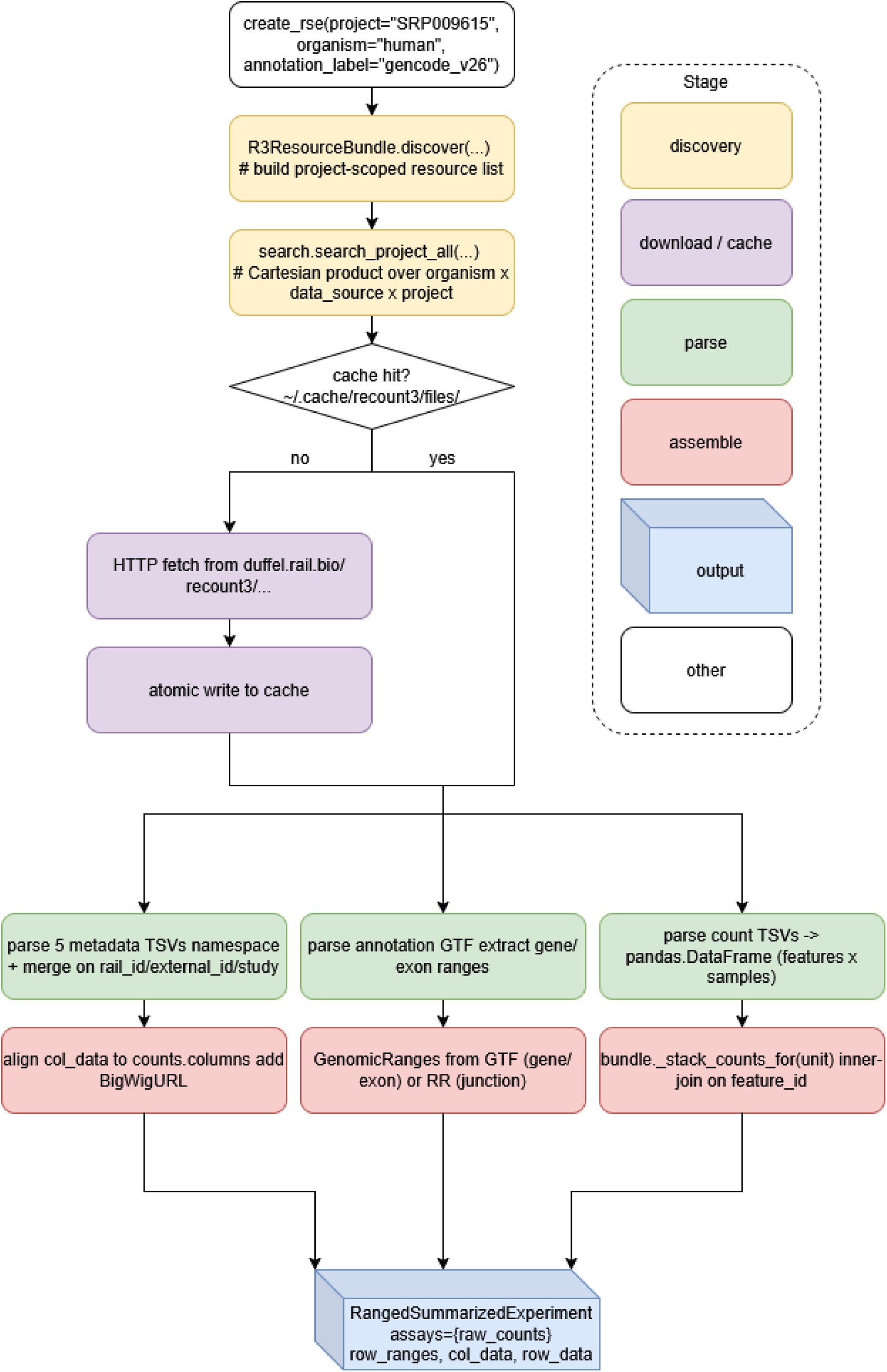
Process flowchart for the construction and instantiation of a RangedSummarizedExperment object.

The recount3.se submodule includes additional utility functions for data and metadata processing. The compute_read_counts function computes approximate read counts from coverage sums using each sample’s average mapped read length. The compute_tpm function calculates transcripts per million (TPM). The compute_scale_factors function computes per-sample scale factors using either the area under the coverage (AUC) or the mapped-reads method and the transform_counts function applies those factors to a count matrix. The is_paired_end function infers paired-end status from the metadata. The expand_sra_attributes function expands encoded SRA attribute strings into distinct metadata columns.

### Command-line interface (CLI)

The package provides a command-line interface (CLI) to support data analysis pipelines and shell scripting. The CLI mirrors the API’s discover, manifest, and materialize workflow through independent subcommands: ids, search, download, bundle, and smoke-test. Figure 4 shows the basic help documentation provided by the CLI.

**Figure 4.**
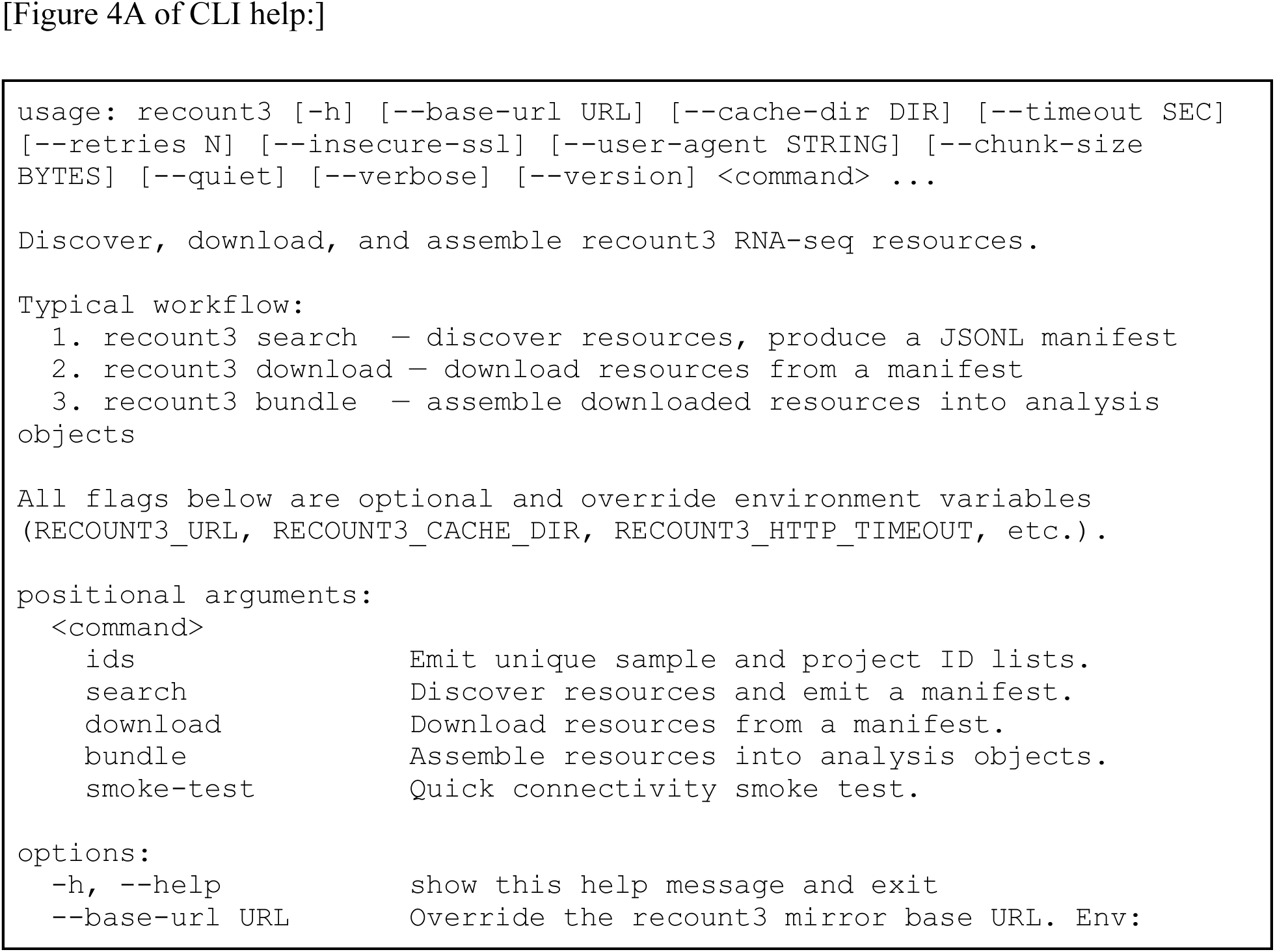
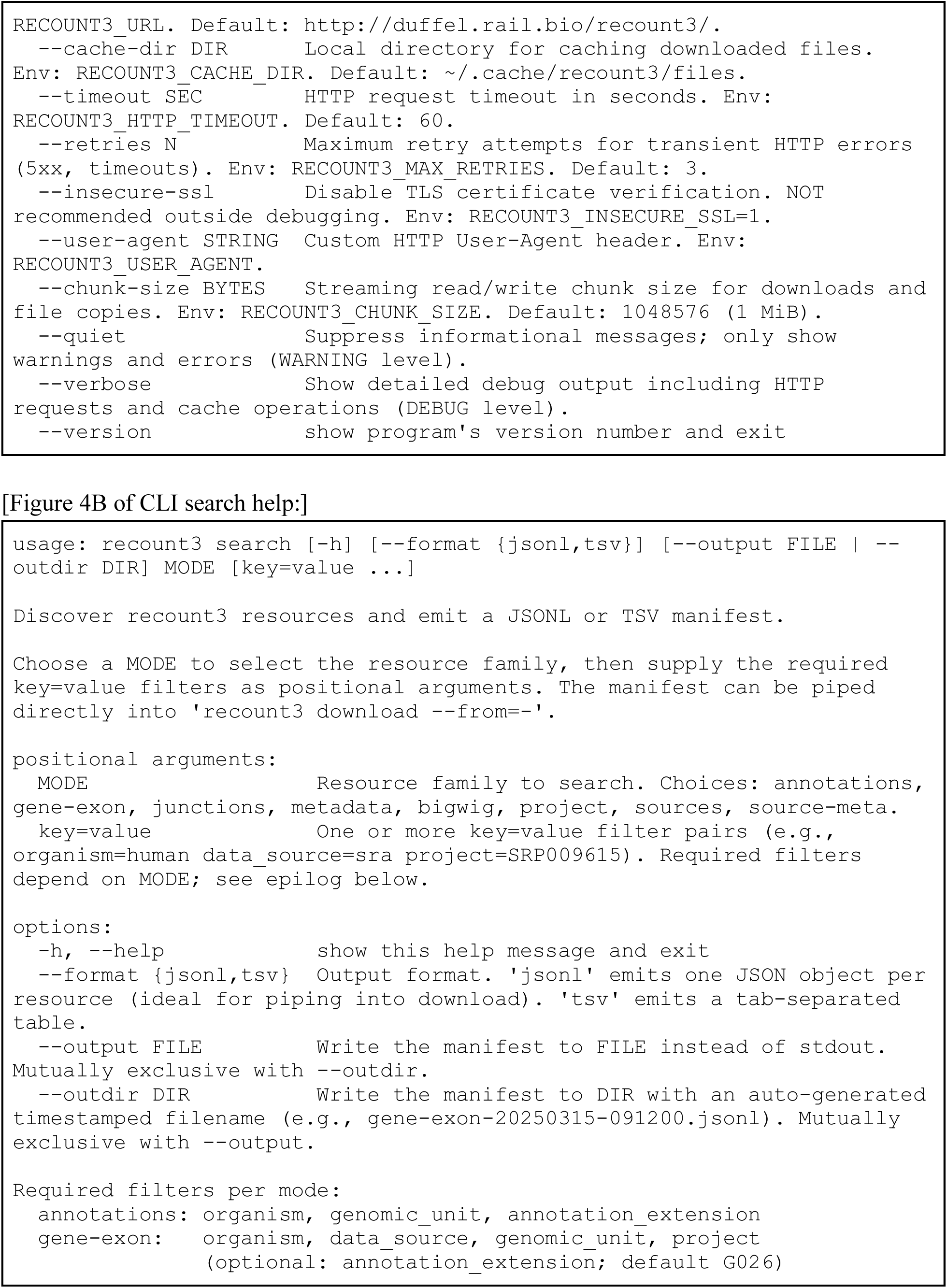
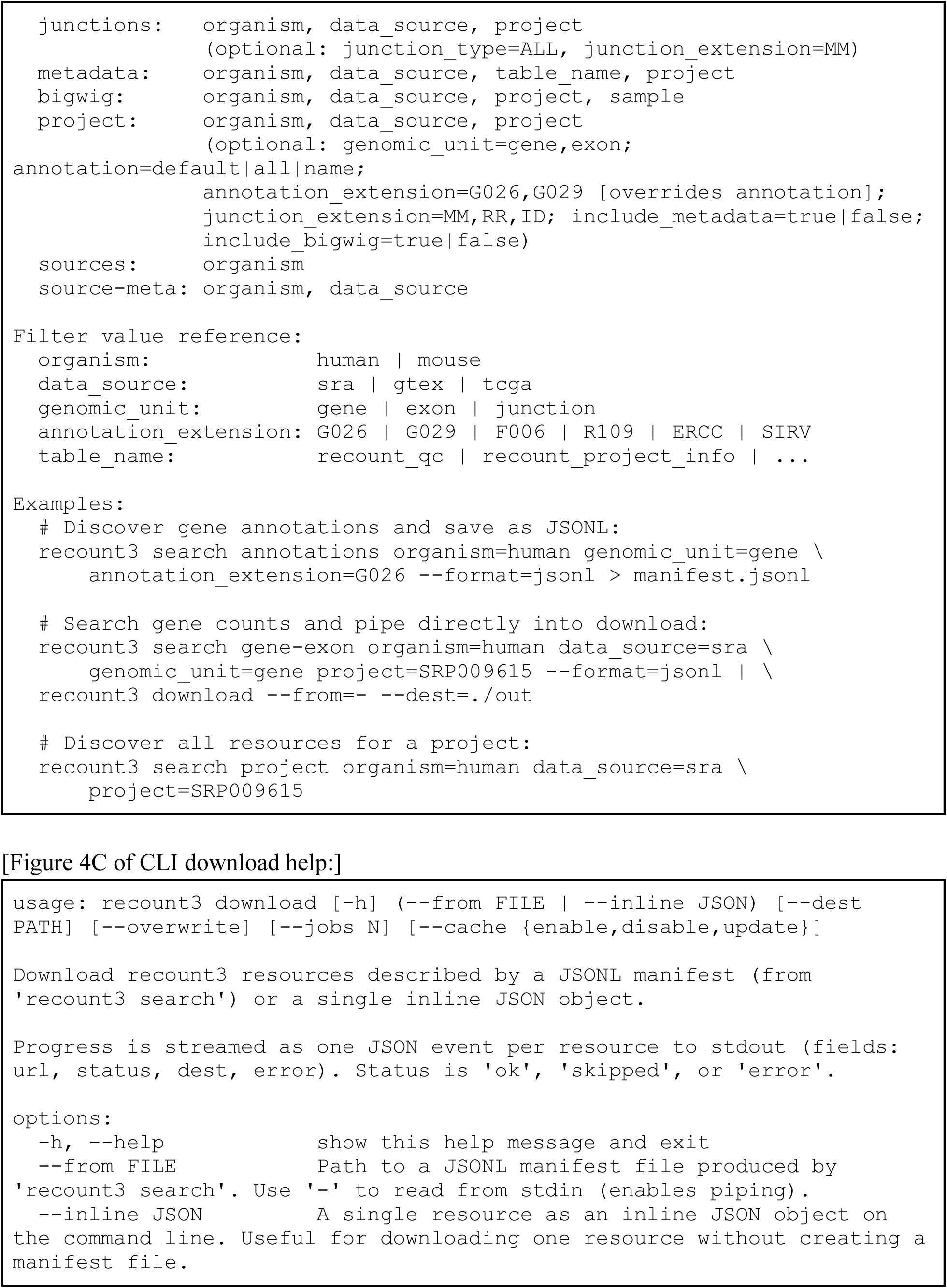
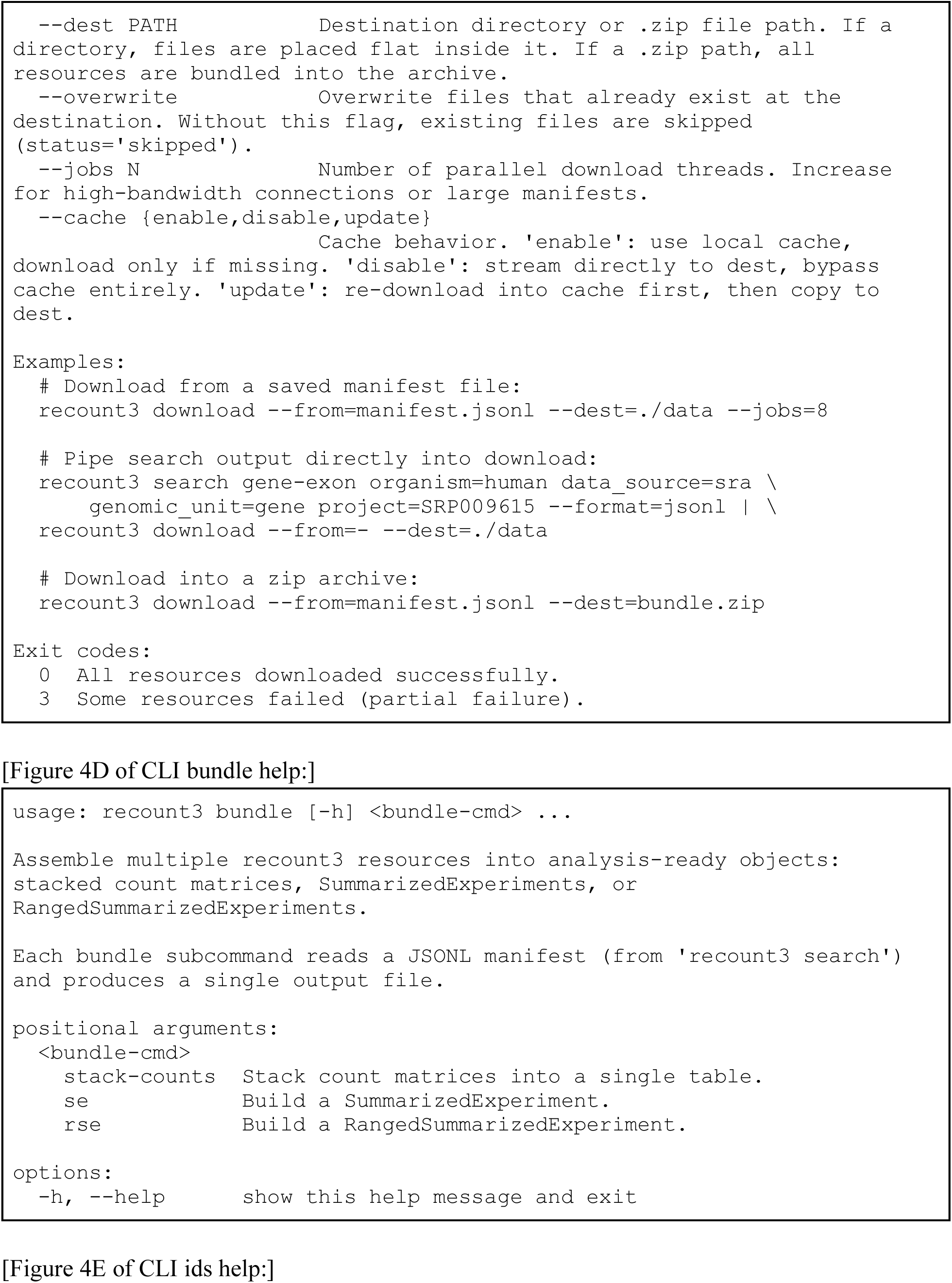
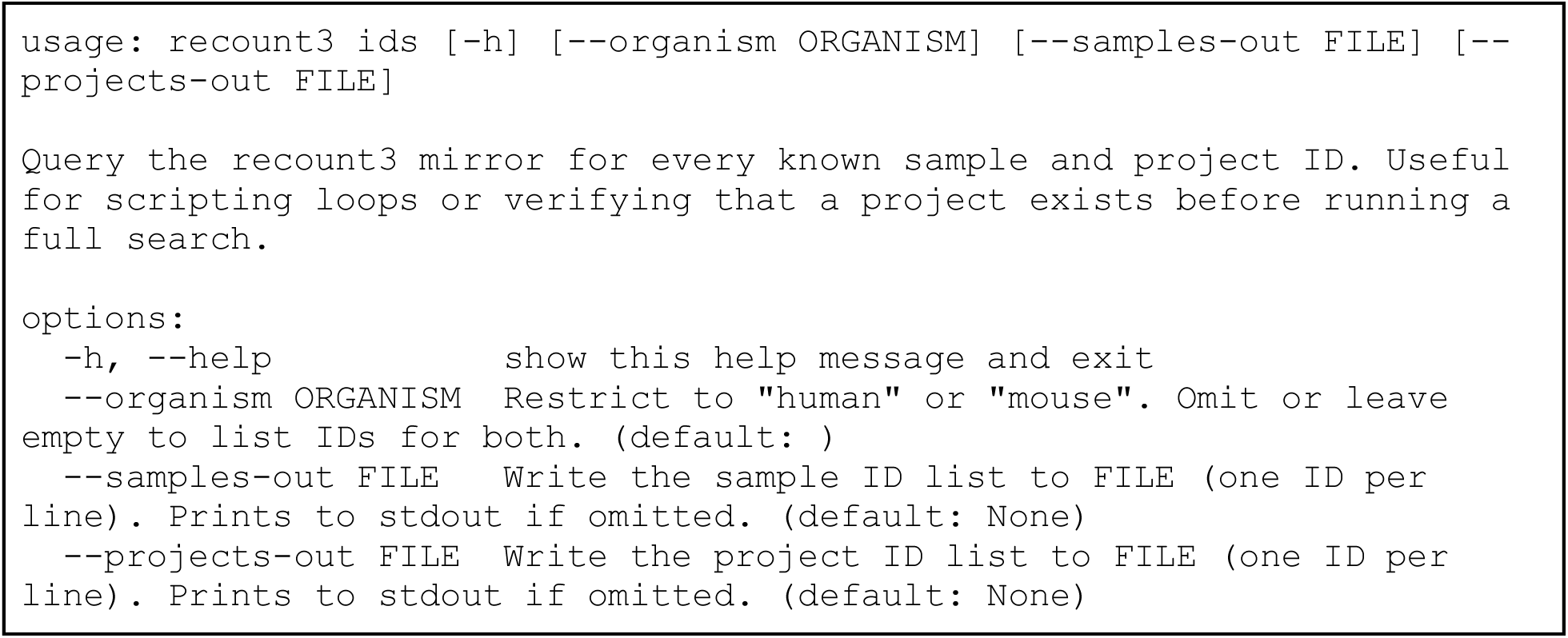
The recount3 CLI help documentation. A) The top-level CLI help documentation. B) The search subcommand help documentation. C) The download subcommand help documentation. D) The bundle subcommand help documentation. E) The ids subcommand help documentation.

**Figure 5.**
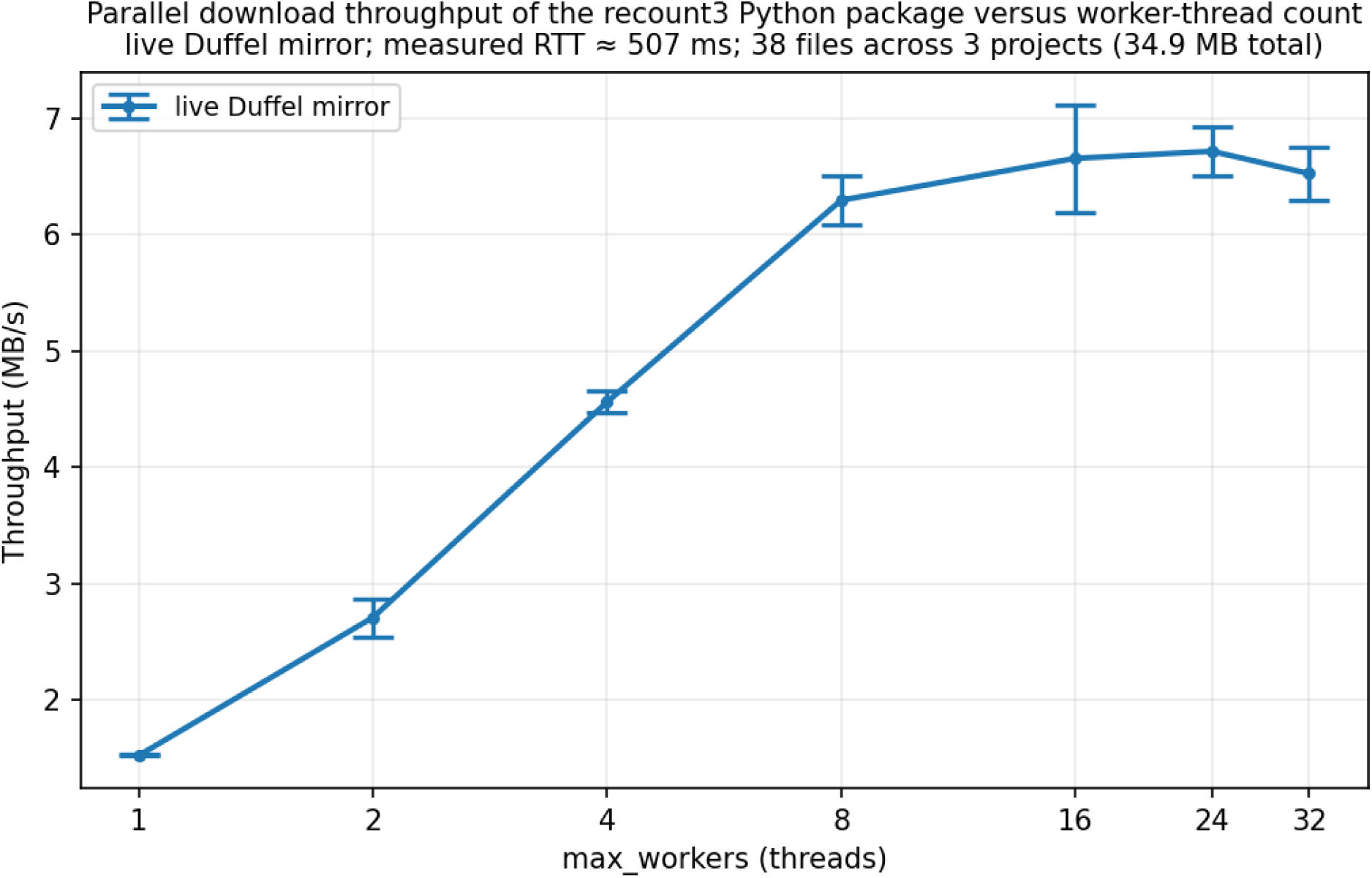
Parallel download benchmark. Figure 5. Graph of number of parallel threads versus download throughput on a 100Mbit/s network connection. The error bars represent the 95% confidence interval estimated from 4 replicate measurements based on a Student’s t distribution.

The search subcommand emits machine-readable JSON Lines (JSONL) or tab-separated values (TSV) manifests, one structured record per resource. JSONL output is designed to be piped directly into the download subcommand, which reads the stream from a file or standard input and fetches the data through a thread pool sized by --jobs. The two subcommands together form a stream-symmetric pipeline (recount3 search … --format=jsonl | recount3 download --from=-) that requires no intermediate file. The download subcommand emits one JSON event per resource to standard output (with status ∈ {ok, skipped, error}), exits with code 0 on full success, 3 on partial failure, and 2 on fatal errors. Destination directories are created automatically (symmetric with the auto-creation already performed for .zip destinations).

The bundle subcommand reads a manifest and produces analysis-ready outputs through three sub-subcommands: stack-counts (TSV, gzip TSV, or Apache Parquet), se and rse (pickled SummarizedExperiment / RangedSummarizedExperiment, with optional AnnData HDF5 output via an anndata dependency). The ids and smoke-test subcommands round out the surface by exposing the full sample and project identifier listings, and by providing a quick connectivity probe for continuous-integration environments. Beyond automated pipelines, the CLI provides a rapid method for data inspection: researchers can preview available projects, query metadata, or check file sizes without launching a Python interpreter, facilitating faster exploration of the repository landscape.

CLI flags mirror the Config dataclass exactly (--base-url, --cache-dir, --timeout, --retries, --insecure-ssl, --user-agent, --chunk-size, --cache=enable|disable|update), and take precedence over environment variables, which themselves take precedence over library defaults.

### Software tool requirements

The recount3 Python package requires Python version 3.10 or higher. Core package dependencies include numpy (≥ 2.0) [9], pandas (≥ 2.2) [10], and scipy (≥ 1.13) [11]. The interoperability functionality requires the optional BiocPy ecosystem packages biocframe (≥ 0.7), summarizedexperiment (≥ 0.6), and genomicranges (≥ 0.8) [13]. BigWig file access requires the pyBigWig library (≥ 0.3.18) [6]. Source code and documentation are distributed through GitHub and GitHub Pages, respectively. Package releases are distributed via the Python Package Index (PyPI) using pyproject.toml packaging (PEP 518).

### Parallel download benchmark

To quantify the benefit of the multithreaded downloading exposed by R3ResourceBundle.download() (and, equivalently, by the CLI download subcommand’s --jobs option), aggregate download throughput was measured as a function of the number of worker threads (max_workers) against the live recount3 “Duffel” mirror.

The benchmark workload was assembled to be representative of small per-project retrievals yet deliberately bounded, so as not to burden the public mirror. Small per-project files, gene and exon count matrices, the per-project metadata tables, and junction sidecar files, were enumerated across three human SRA projects (SRP009615, SRP014565, SRP096765) using the package’s own offline search_* functions, and each candidate was pre-screened with a header-only HTTP request so that only files that already existed and were at most 8 MB were retained, up to a 50 MB aggregate budget. This procedure yielded 38 files totalling 34.9 MB. Downloads were performed with caching disabled (cache_mode=“disable”) so that every trial constituted a network transfer rather than a cache hit; transferred files were written to a scratch directory on local storage and deleted between trials, and the persistent on-disk cache was neither read nor written.

For each thread count in {1, 2, 4, 8, 16, 24, 32}, a single warm-up download was discarded, and four timed downloads were recorded, with trial order interleaved across thread counts under a fixed pseudorandom seed to average out slow drift in network conditions. Throughput was computed as the total number of bytes transferred divided by wall-clock time, and parallel speedup as the ratio of single-thread to k-thread completion time; error bars denote 95% confidence intervals estimated from four replicates based on a Student’s t distribution.

Measurements were collected with an Intel Xeon E5-1650 v4 processor (6 physical cores, 12 hardware threads via simultaneous multithreading) and 32 GB of DDR4-2400 ECC memory (4 × 8 GB) in a quad-channel configuration, of which 16 GB was made available to the Linux guest. Transient download files were written to an SK Hynix Gold P31 1 TB NVMe solid-state drive, ensuring that local storage was not a throughput bottleneck. The host’s network access was provided over a 100 Mbit/s Ethernet link. The software environment was Debian GNU/Linux 13 (trixie) running under the Windows Subsystem for Linux 2 (WSL2) on Windows 10, with Python 3.14.

## Declarations

### Ethics approval and consent to participate

Not applicable

### Consent for publication

Not applicable

### Availability of data and materials

The recount3 Python package is available on GitHub: https://github.com/MoseleyBioinformaticsLab/recount3

On the Python Package Index (PyPI) https://pypi.org/project/recount3/

With full end-user documentation: https://moseleybioinformaticslab.github.io/recount3/

### Authors’ contributions

Conceptualization, HNBM and RMF; methodology, HNBM, AA, and RMF; software, AA; validation, AA and HNBM; formal analysis, AA; investigation, AA; resources, HNBM; writing—original draft preparation, AA and HNBM; writing—review and editing, HNBM, AA, and RMF; visualization, AA; supervision, HNBM; project administration, HNBM; funding acquisition, HNBM. All authors have read and agreed to the published version of the manuscript.

## Acknowledgements

We would like to thank Mr. Erik Huckvale for suggestions and help with generating UML diagrams.

## Supplemental Material for

**Supplemental Table 1.**
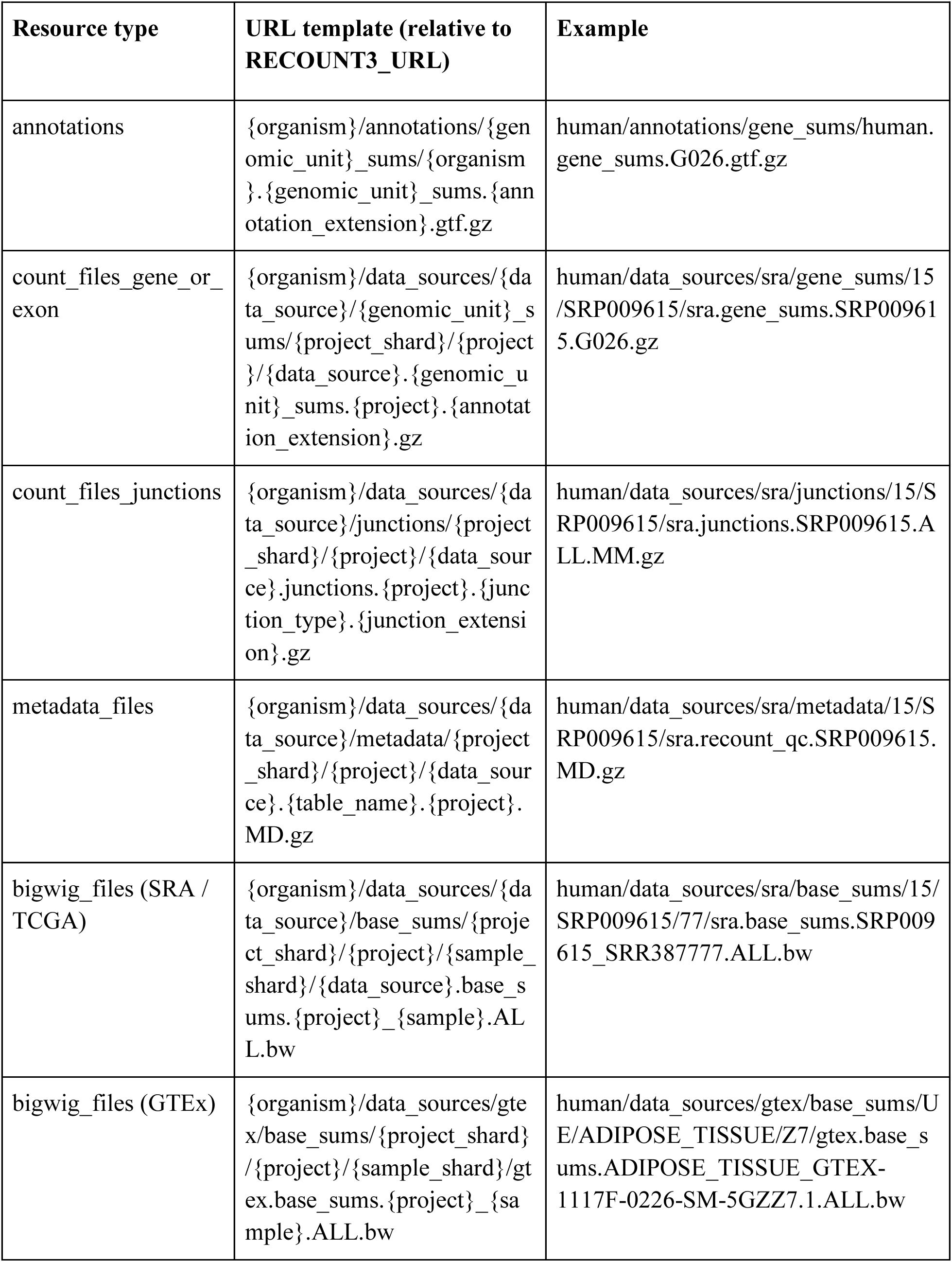
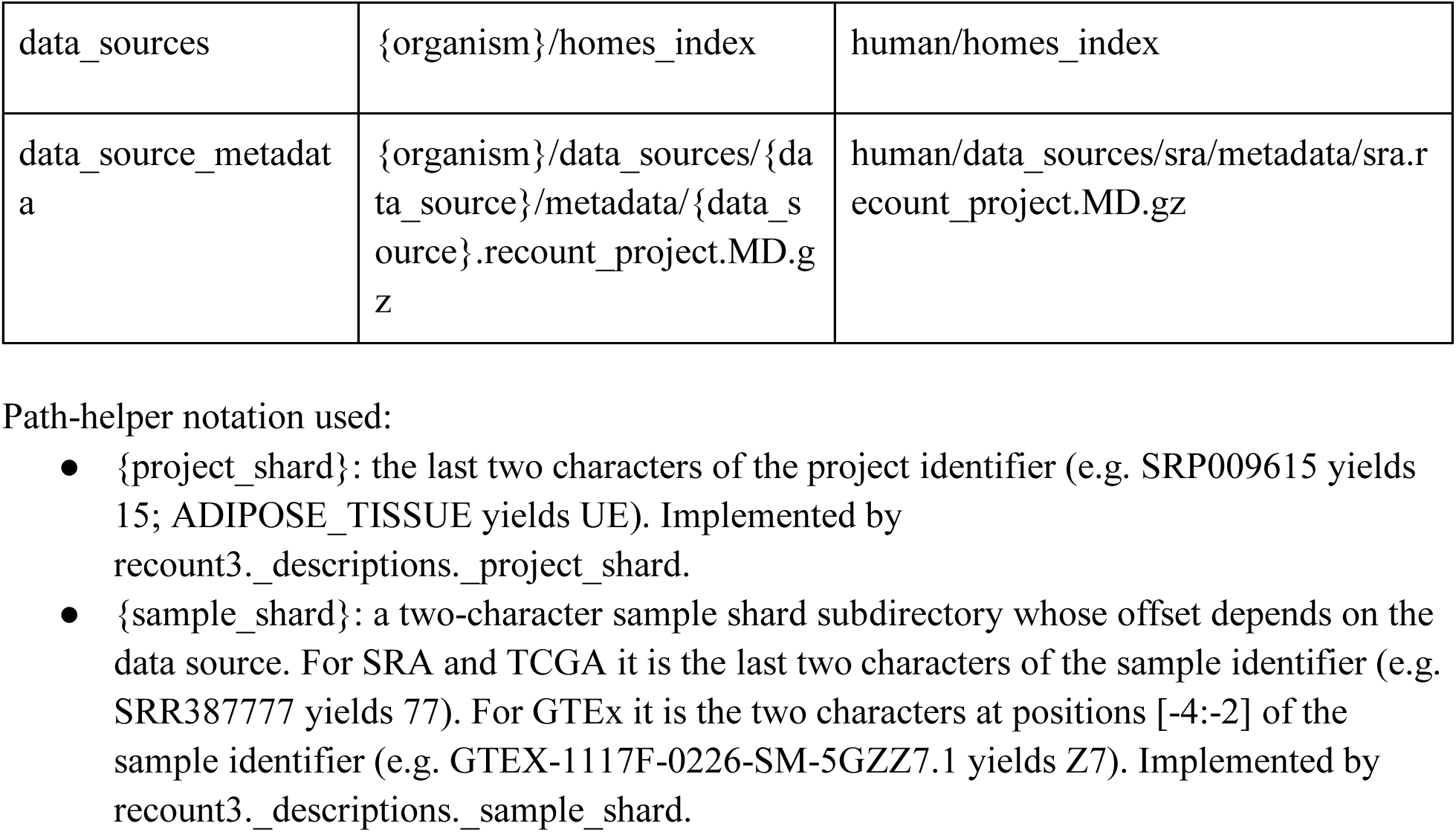
URL templates for downloading recount3 files.

Path-helper notation used:

● {project_shard}: the last two characters of the project identifier (e.g. SRP009615 yields 15; ADIPOSE_TISSUE yields UE). Implemented by recount3._descriptions._project_shard.
● {sample_shard}: a two-character sample shard subdirectory whose offset depends on the data source. For SRA and TCGA it is the last two characters of the sample identifier (e.g. SRR387777 yields 77). For GTEx it is the two characters at positions [-4:-2] of the sample identifier (e.g. GTEX-1117F-0226-SM-5GZZ7.1 yields Z7). Implemented by recount3._descriptions._sample_shard.

## References

1. Wang Z, Gerstein M, Snyder M: RNA-Seq: a revolutionary tool for transcriptomics. Nature reviews genetics 2009, 10(1):57–63.

2. Katz K, Shutov O, Lapoint R, Kimelman M, Brister JR, O’Sullivan C: The Sequence Read Archive: a decade more of explosive growth. Nucleic acids research 2022, 50(D1):D387–D390.

3. Wilks C, Zheng SC, Chen FY, Charles R, Solomon B, Ling JP, Imada EL, Zhang D, Joseph L, Leek JT: recount3: summaries and queries for large-scale RNA-seq expression and splicing. Genome biology 2021, 22(1):1–40.

4. Lonsdale J, Thomas J, Salvatore M, Phillips R, Lo E, Shad S, Hasz R, Walters G, Garcia F, Young N: The genotype-tissue expression (GTEx) project. Nature genetics 2013, 45(6):580–585.

5. Weinstein JN, Collisson EA, Mills GB, Shaw KR, Ozenberger BA, Ellrott K, Shmulevich I, Sander C, Stuart JM: The cancer genome atlas pan-cancer analysis project. Nature genetics 2013, 45(10):1113–1120.

6. Ryan D, Grüning B, Ramirez F: pyBigWig 0.2. 4. *Zenodo* 2016.

7. JSON Lines [https://jsonlines.org/]

8. Vohra D: Apache parquet. In: Practical Hadoop Ecosystem: A Definitive Guide to Hadoop-Related Frameworks and Tools. Springer; 2016: 325–335.

9. Harris CR, Millman KJ, Van Der Walt SJ, Gommers R, Virtanen P, Cournapeau D, Wieser E, Taylor J, Berg S, Smith NJ: Array programming with NumPy. Nature 2020, 585(7825):357–362.

10. McKinney W: pandas: a foundational Python library for data analysis and statistics. Python for high performance and scientific computing 2011, 14(9):1–9.

11. Virtanen P, Gommers R, Oliphant TE, Haberland M, Reddy T, Cournapeau D, Burovski E, Peterson P, Weckesser W, Bright J: SciPy 1.0: fundamental algorithms for scientific computing in Python. Nature methods 2020, 17(3):261–272.

12. Virshup I, Rybakov S, Theis FJ, Angerer P, Wolf FA: anndata: Access and store annotated data matrices. Journal of Open Source Software 2024, 9(101):4371.

13. BiocPy: Facilitate Bioconductor Workflows in Python [https://github.com/biocpy]

14. Gumienny R, van Heeringen S, Bismeijer T, Ramdhani H: GEOparse: Python library to access gene expression omnibus database (GEO). In.: Python Package Index; 2019.

15. Lachmann A, Torre D, Keenan AB, Jagodnik KM, Lee HJ, Wang L, Silverstein MC, Ma’ayan A: Massive mining of publicly available RNA-seq data from human and mouse. Nature communications 2018, 9(1):1366.

16. Köster J, Rahmann S: Snakemake—a scalable bioinformatics workflow engine. Bioinformatics 2012, 28(19):2520–2522.

17. Di Tommaso P, Chatzou M, Floden EW, Barja PP, Palumbo E, Notredame C: Nextflow enables reproducible computational workflows. Nature biotechnology 2017, 35(4):316–319.

18. Fielding RT: Architectural styles and the design of network-based software architectures: University of California, Irvine; 2000.

19. Hunt J: Data Classes. In: Advanced Guide to Python 3 Programming. Springer; 2023: 33–47.

20. Registry Pattern [https://www.geeksforgeeks.org/system-design/registry-pattern/]

21. Gamma E, Helm R, Johnson R, Vlissides J: Design patterns: Abstraction and reuse of object-oriented design. In: European conference on object-oriented programming: 1993: Springer; 1993: 406–431.

22. Boisvert RF, Pozo R, Remington K, Barrett RF, Dongarra JJ: Matrix Market: a web resource for test matrix collections. In: Quality of Numerical Software: Assessment and Enhancement. Springer; 1997: 125–137.

